# Single amino acid mutations in histone H3.3 illuminate the functional significance of H3K4 methylation in plants

**DOI:** 10.1101/2024.12.02.626315

**Authors:** Mande Xue, Lijun Ma, Xiaoyi Li, Huairen Zhang, Qian Liu, Danhua Jiang

**Affiliations:** Key Laboratory of Seed Innovation, Institute of Genetics and Developmental Biology, Chinese Academy of Sciences, Beijing, China; University of Chinese Academy of Sciences, Beijing, China; Temasek Life Sciences Laboratory, 1 Research Link, National University of Singapore, Singapore

## Abstract

Although numerous data have revealed correlations between histone modifications and chromatin activities such as transcription, proofs of their causal importance in gene expression regulation remain limited. Sequence variants within each histone family expand chromatin diversity and may carry specific modifications, further raising the question of how histone modifications coordinate with different variants. Here, we investigate the regulatory role of lysine 4 (K4) of *Arabidopsis* histone H3 by point mutating K4 in two major H3 variants, H3.1 and H3.3. K4 is essential for the function of H3.3 but not H3.1 in plant development. H3K4 methylation levels decrease drastically upon K4 mutation in H3.3, and the associated transcriptome changes are similar to those observed in a mutant lacking SDG2, a major enzyme responsible for depositing H3K4 trimethylation (H3K4me3). Moreover, H3.3K4 and SDG2 are required for de novo gene activation and RNA Pol II elongation. H3K4 methylation is preferentially accumulated on H3.3 compared to H3.1, likely due to the close association of the H3.3 deposition and H3K4 methylation machineries. Furthermore, we reveal the diverse impacts of K4 nearby residue mutations on H3K4 methylation and H3.3 function. Collectively, these findings suggest that H3.3 serves as a critical substrate for H3K4 methylation, which is important for gene expression regulation. In addition, our work highlights the potential of using plants as a platform to directly investigate the importance of histone amino acids and their modifications.

## Introduction

Histone modification is widely considered as a crucial mechanism for controlling eukaryotic genome activity. Various modifications, such as acetylation, methylation, phosphorylation, and ubiquitination, have been detected on histone residues ^1,2^. Histone acetylation involves the addition of a negative charged acetyl group to the histone lysine (K) residues, neutralizing their positive charge and weakening the interaction between DNA and histone proteins. As a result, histone acetylation is suggested to promote an open chromatin structure permissive for transcription ^3,4^. Compared with acetylation, methylation on histone K or arginine (R) residues is associated with a more varied impact on chromatin. For instance, at protein-coding genes, the presence of histone H3K27 trimethylation (H3K27me3) is correlated with transcriptional repression, while active genes often show an enrichment of H3K4 and H3K36 methylation ^1,2^. Most of the studies addressing the significance of histone modifications have relied on the disruption of enzymes catalysing these modifications. However, non-histone substrates or enzymatic independent activities have been reported for these enzymes ^5-10^. Therefore, despite the well-established correlations between histone modifications and transcriptional activity, the direct requirement of these modifications on transcription remains largely unclear. Recent studies have employed a point mutation approach by specifically mutating the modified residues such as H3K27, which suggest the importance of H3K27 methylation in modulating transcription ^11-15^.

H3K4 trimethylation (H3K4me3) is a conserved, permissive histone modification found from yeast to animals and plants. It normally accumulates around the transcription start site of active genes. In yeast, H3K4me3 is catalysed by the COMPASS complex (complex of proteins associating with Set1), comprising the histone methyltransferase Set1 and several core structural components such as SWD1, SWD2, SWD3, and BRE2 ^16,17^. COMPASS-like complexes have been identified in animals and plants and shown to play conserved functions in depositing H3K4 methylation ^18-22^. In *Arabidopsis*, RBL, S2Lb, WDR5a, and ASH2R are homologs of SWD1, SWD2, SWD3, and BRE2 respectively ^18-20^, and several SET domain-containing proteins are responsible for catalysing H3K4 methylation ^23-29^. For example, *Arabidopsis* SET DOMAIN GROUP 2 (SDG2) is identified as a major H3K4 methyltransferase broadly contributing to H3K4me3. Although it can catalyse all three forms of H3K4 methylation *in vitro*, loss of SDG2 only causes a strong reduction in H3K4me3 but not H3K4 mono- and dimethylation (H3K4me1 and H3K4me2) ^23,26^. This probably is due to the functional redundance of H3K4 methyltransferases in depositing H3K4me1 and H3K4me2 ^25,30^. Loss of SDG2 or other subunits of the COMPASS-like complex induces gene misregulation and severe developmental defects, including dwarfism, early flowering, and infertility ^18,20,23,25-27,29^. However, H3K4 methyltransferases have been reported to methylate non-histone substrates such as Borealin in the chromosome passenger complex ^31^, and COMPASS-like complexes have been suggested to also regulate transcription or other cellular activities independent of its catalytic roles ^32-36^. So far, there is a lack of studies addressing the causal importance of H3K4 methylation in transcriptional regulation and development, particularly in plants.

Besides H3K4me3, another player in chromatin regulation that is closely linked with active transcription is the histone variant H3.3 ^37-39^. Histone variants are related but functional diversified members in the same histone family ^40,41^. In both animals and plants, H3.3 differs from another major H3 variant H3.1/H3.2 by only 4-5 amino acids ^42,43^. Yet compared to H3.1, H3.3 shows greater enrichment at actively transcribed regions ^37-39^. In addition, H3.1 and H3.3 have distinct deposition modes. H3.1 is incorporated into chromatin by the CHROMATIN ASSEMBLY FACTOR-1 (CAF1) complex in a DNA replication-dependent manner ^11,44^, while H3.3 is replication-independently incorporated by the HISTONE REGULATORY HOMOLOG A (HIRA) complex and ALPHA THALASSEMIA MENTAL RETARDATION SYNDROME X-LINKED (ATRX)-DAXX ^37,45-47^. The HIRA complex is evolutionarily conserved and consists of HIRA, UBN1/2, and CABIN1 ^48,49^. In mammalian cells, UBN1/2 preferentially interact with H3.3, thereby conferring specificity for H3.3 binding by the HIRA complex ^50,51^.

Although much progress has been made on the chromatin localization and deposition mechanisms of H3.3, its molecular function remains elusive. In *Drosophila*, loss of H3.3 does not affect viability and only causes male sterile ^52,53^, while in *Xenopus* and mouse, H3.3 is essential for embryo development ^54,55^. Knockdown of H3.3 in *Arabidopsis* leads to leaf serration, early flowering, reduced fertility and impaired high temperature response ^56-58^, and complete deletion of H3.3 causes severe defects in germination and post-embryonic development ^59^. H3.3 may regulate chromatin activity through modulating histone modifications. Compared with H3.1, H3.3 contains a specific serine (S) 31 (in animals) or threonine (T) 31 (in plants) residue at its N-terminus ^42,43^. In *Xenopus* embryo, mimicking phosphorylation at S31 stimulates acetylation at K27, which is permissive for transcription ^55^. In mouse cells, stimulation induces phosphorylation at H3.3S31, which attracts a histone methyltransferase SETD2 for the deposition of H3K36me3 ^60^. *Arabidopsis* H3.3 is immune to the repressive H3K27me1 catalysed by the plant-specific methyltransferases, *ARABIDOPSIS* TRITHORAX-RELATED PROTEINS 5 (ATXR5) and ATXR6, as the H3.3T31 residue inhibits their activity ^61^. Thus, dissecting the role of individual amino acids in H3.3 and their impacts on histone modifications is crucial for obtaining a deeper understanding of the H3.3 function.

In this study, we investigate the requirement of the K4 residue in H3.1 or H3.3 by complementing their respective mutants with K4 mutated versions. Changes of K4 to other amino acids impair the function of H3.3 but not H3.1. The H3.3K4 mutation induces a strong reduction in global H3K4 methylation levels, which is associated with drastic transcriptomic changes and de novo gene activation defects that are similar to those in a *sdg2* mutant. We show that H3K4 methylation is predominantly deposited on H3.3, probably because of the close association of the COMPASS-like complex with the H3.3 chaperone HIRA complex. In addition, our single amino acid mutation strategy uncovers the significance of H3.3K4 neighbouring residues and the varied impacts of their mutations on H3K4 methylation and the function of H3.3. Together, these results suggest the biological importance of H3K4 methylation and pinpoint the critical requirement of H3.3 for its deposition.

## Results

### The H3.3K4 residue is essential for plant development

Several lysine residues in H3 carry modifications and they are identical in H3.1 and H3.3. In this study, we focused on the N-terminal K4, K9, and K27 residues and investigated their importance for H3.1/H3.3 (Figure 1a). We introduced mutations into *HTR5* (an H3.3 coding gene) or *HTR13* (an H3.1 coding gene) to switch each of these lysine residues to alanine (A), and used these mutated forms for complementation tests. Like previously reported ^11^, the K4 and K9 but not K27 mutated H3.1 successfully complemented the inflorescence enlargement phenotype of an *h3.1* knockdown mutant (*h3.1kd*) (Supplemental Figure 1), indicating that the K4 and K9 residues in H3.1 are not essential for its function in plant development. For H3.3, all its mutated forms were able to rescue the strong germination defects of an *h3.3* complete knockout mutant (*h3.3ko*) lacking all three H3.3 coding genes *HTR4*, *HTR5*, and *HTR8* ^59^, with K4 mutated H3.3 showing slightly reduced capability (Figure 1b and Supplemental Figure 2a). After germination, while the non-mutated and K9/K27 mutated H3.3 could fully rescue the *h3.3ko* phenotypes, resulting in normally developed plants similar to the wild type (WT) Columbia (Col) (Figure 1c), all independent T3 transgenic lines carrying the K4 mutated H3.3 displayed strong developmental abnormities (phenotypes of three representative lines are shown), including smaller plants (Figure 1c and 1d and Supplemental Figure 2b and 2c), early flowering (Figure 1e and Supplemental Figure 2d), abnormally developed flowers (Figure 1f), and complete sterility (Figure 1g and Supplemental Figure 2e).

**Figure 1.**
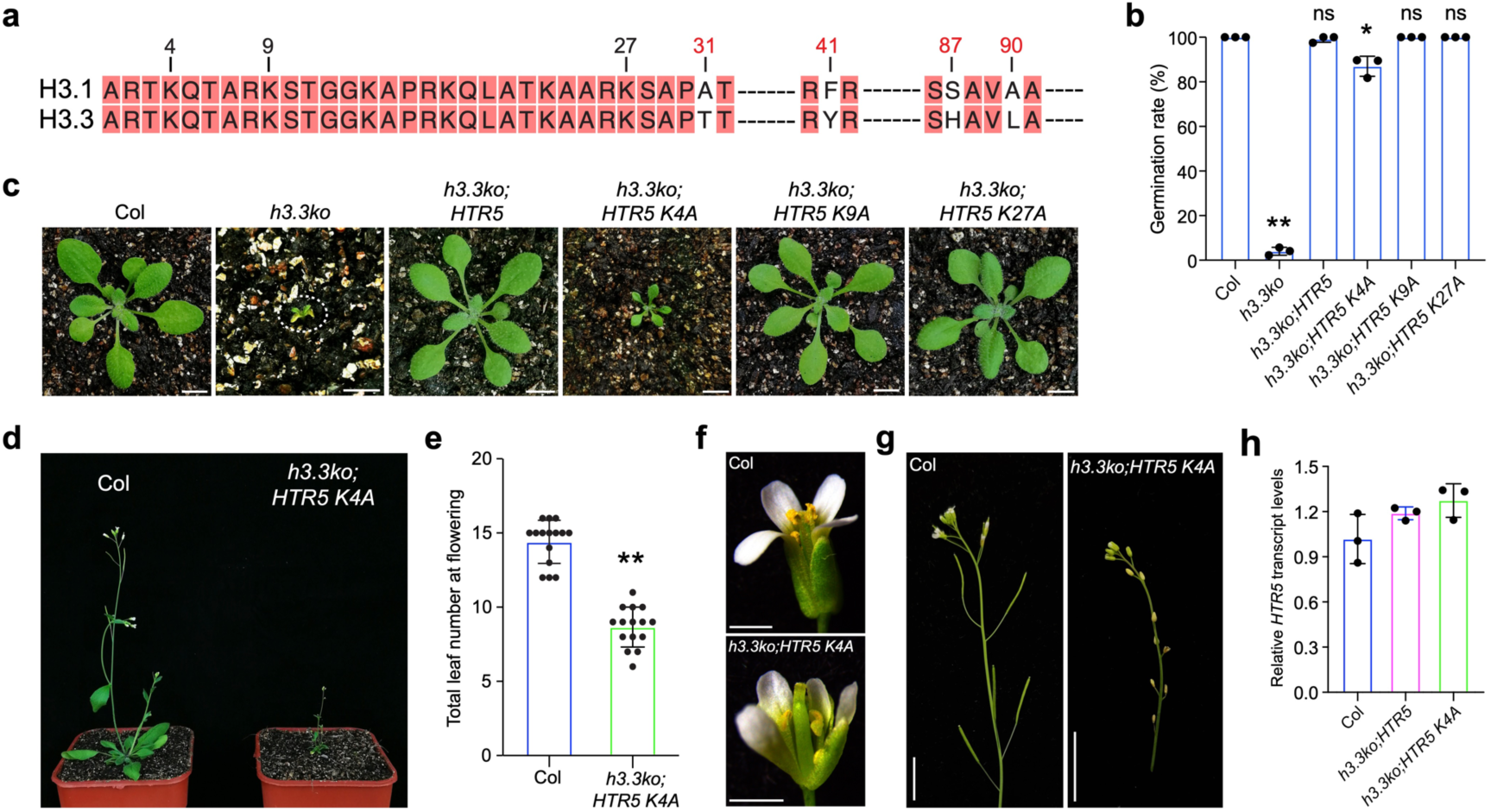
H3.3K4 residue is essential for the H3.3 function in plant development. **a.** Sequence alignment of *Arabidopsis* H3.1 and H3.3. Numbers indicate the positions of lysine residues mutated in this study and highlight the residues that differentiate H3.1 and H3.3. **b.** Seed germination rates of the indicated lines after imbibed for 7 days. Values are means ± SD of three biological replicates. At least 32 seeds were analysed per replicate. Statistical significance relative to Col was determined by two-tailed Student’s *t*-test (*, *P* < 0.05; **, *P* < 0.01; ns, not significant). **c.** Developmental phenotypes of the indicated lines at the vegetative stage. Scale bars, 1 cm. **d.** Developmental phenotypes of Col and *h3.3ko;HTR5 K4A* at the reproductive stage. **e.** Total number of primary rosette and cauline leaves at flowering for Col and *h3.3ko;HTR5 K4A*. 15 plants were scored for each line. Values are means ± SD. The significance of differences was determined by two-tailed Student’s *t*-test (**, *P* < 0.01). **f and g.** Flower (f) and silique (g) developmental phenotypes of Col and *h3.3ko;HTR5 K4A*. Scale bars, 1 mm (f) or 1 cm (g). **h.** Relative *HTR5* transcript levels in the indicated lines determined by RT-qPCR. Values are means ± SD of three biological replicates.

**Figure 2.**
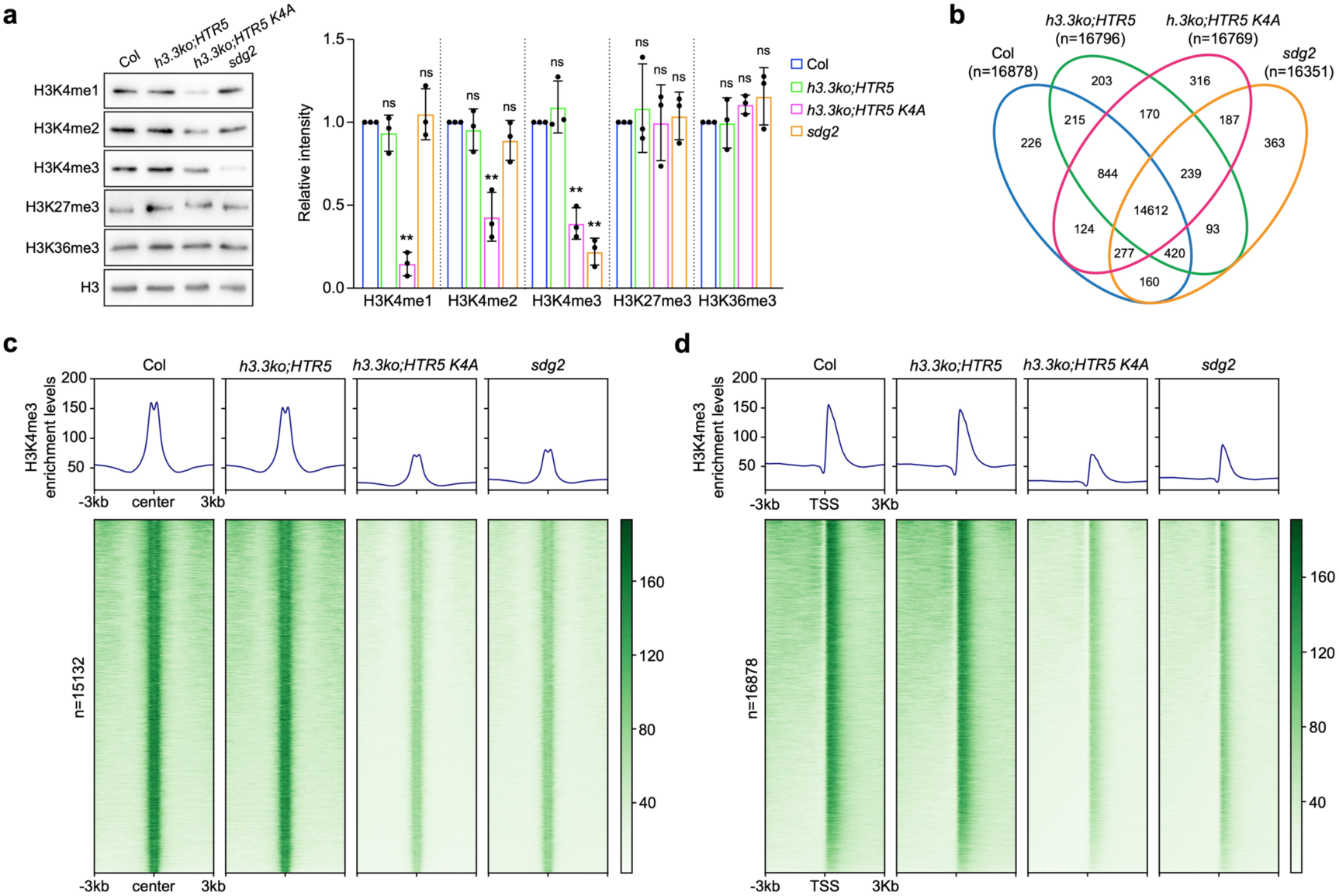
The H3.3K4 is essential for global H3K4 methylation. **a.** Histone modification levels in Col, *h3.3ko;HTR5*, *h3.3ko;HTR5 K4A* and *sdg2* determined by western blotting. H3 was employed as a loading control. The bar chart represents quantification of western blot signals from three biological replicates (Figure 2a and Supplemental Figure 4a). Values are means ± SD. Statistical significance relative to Col was determined by two-tailed Student’s *t*-test (**, *P* < 0.01; ns, not significant). **b.** Venn diagram of H3K4me3-enriched genes in Col, *h3.3ko;HTR5*, *h3.3ko;HTR5 K4A* and *sdg2* determined by ChIP-seq. **c.** Metaplot and heatmap of H3K4me3 ChIP-seq signals in Col, *h3.3ko;HTR5*, *h3.3ko;HTR5 K4A* and *sdg2* over H3K4me3-enriched peaks in WT Col. **d.** Metaplot and heatmap of H3K4me3 ChIP-seq signals in Col, *h3.3ko;HTR5*, *h3.3ko;HTR5 K4A* and *sdg2* over the TSS of H3K4me3-enriched genes in WT Col.

Several mutations of H3 lysine residues, such as K4 to methionine (K4M), K9M, K27M and K36M, can induce gain-of-function effects, likely by dominantly inhibiting the activity of their corresponding histone methyltransferases ^62-67^. We excluded this possibility for the H3.3 K4A, as it did not cause any defects in the *h3.3ko* heterozygous (*htr4/htr4;htr5/htr5;htr8/+*) background (Supplemental Figure 2f). Lysine and arginine (R) are positively charged amino acids. To address the charge-neutralization effect caused by the K4A mutation, we introduced H3.3 K4R into the *h3.3ko* and observed similar developmental defects as those induced by the K4A mutation (Supplemental Figure 2g and 2h), suggesting that the modifications, rather than the positive charge on the H3.3K4 residue, are essential for plant development. K4 can be methylated or acetylated, and the acetylated lysine can be mimicked by lysine to glutamine (Q) substitution ^68,69^. We found that expressing H3.3 K4Q also failed to rescue the *h3.3ko* mutant phenotypes (Supplemental Figure 2i and 2j). This suggests that the methylation on K4 is likely required for the function of H3.3. Since all *h3.3ko* rescue lines expressing H3.3 K4A showed similar phenotypes, we selected line #1 for subsequent studies and referred to it as *h3.3ko;HTR5 K4A*. The expression levels of *HTR5* in this line were comparable to those in WT (Figure 1h).

### The K4 residue in H3.3 is not required for its chromatin distribution

Mutations in H3 proteins may alter their chromatin incorporation ^70,71^, we thus examined whether the K4A mutation affects H3.3 distribution. Due to the lack of a plant H3.3 specific antibody, green fluorescent protein (GFP)-fused H3.3 (HTR5) has been used to examine its genomic localization ^39,58,59^. Two independent transgenic lines expressing HTR5-GFP or HTR5 K4A-GFP under the *HTR5* endogenous promoter were examined. The K4-mutated H3.3 localized normally in the nuclei, showing that its subcellular localization is not affected by the mutation (Supplemental Figure 3a). To investigate the global chromatin localization of H3.3 K4A, we performed immunofluorescence staining with extracted leaf nuclei. Like H3.3, H3.3 K4A was localized at euchromatin but not at heterochromatic regions marked by the histone variant H2A.W (Supplemental Figure 3b) ^72^. Thus, the global H3.3 landscape at chromatin remained unchanged regardless of the K4 mutation.

We further performed chromatin immunoprecipitation sequencing (ChIP-seq) to examine the genome-wide localization of H3.3 and H3.3 K4A. Overall, the ChIP-seq profiles of H3.3 and H3.3 K4A were strongly correlated (Supplemental Figure 3c), and both were mainly accumulated at euchromatic regions (Supplemental Figure 3d). In vegetative tissues, H3.3 is mainly enriched at genic regions, and its accumulation levels are positively correlated with gene expression activity ^38,39^. H3.3 K4A showed similar genic distribution patterns to H3.3 (Supplemental Figure 3e). Together, these results suggest that the K4 mutation in H3.3 does not affect its chromatin incorporation.

### H3.3K4 mutation strongly impairs global H3K4 methylation levels

To elucidate the role of the H3.3K4 residue in histone modifications, we examined global levels of several histone modifications by immunoblotting. SDG2 is a major H3K4 methyltransferase responsible for H3K4me3 in *Arabidopsis*, and loss of SDG2 also leads to smaller plants and sterility, similar to the phenotypes of *h3.3ko;HTR5 K4A* ^23,26^. Thus, we included the *sdg2* mutant for comparison in our analysis. Compared with WT and *h3.3ko;HTR5*, all three forms of H3K4 methylation showed reduced levels in *h3.3ko;HTR5 K4A*, while in *sdg2*, only levels of H3K4me3 were significantly reduced. In addition, global levels of other euchromatic modifications, H3K27me3 and H3K36me3, were not altered in *h3.3ko;HTR5 K4A* and *sdg2* (Figure 2a and Supplemental Figure 4a).

H3K4me3 is a well-characterized histone modification linked with active gene transcription. Hence, we focused on it and examined its genome-wide accumulation in *h3.3ko;HTR5 K4A* and *sdg2* by ChIP-seq. Due to the expected global reduction in H3K4me3, we included a spike-in reference (human HEK293 chromatin) in the ChIP-seq experiment ^73^. The number of H3K4me3-enriched peaks and genes were similar in *h3.3ko;HTR5 K4A* and *sdg2* compared with WT and *h3.3ko;HTR5* (Supplemental Figure 4b), and the majority of H3K4me3-enriched genes were still shared among them (Figure 2b), suggesting that the H3K4me3 distribution patterns are maintained in *h3.3ko;HTR5 K4A* and *sdg2*. However, at H3K4me3-enriched peaks in WT, enrichment levels were drastically reduced in *h3.3ko;HTR5 K4A* and *sdg2* compared with WT and *h3.3ko;HTR5* (Figure 2c). Similarly, at H3K4me3-enriched genes, H3K4me3 levels were reduced in *h3.3ko;HTR5 K4A* and *sdg2*, despite they still peaked around the transcription start site (TSS) (Figure 2d). Together, these results demonstrate that loss of the K4 residue in H3.3 compromises global H3K4 methylation levels.

It was previously reported that H3.3 promotes gene body DNA methylation, particularly CG methylation ^56,59^. To determine whether this function of H3.3 relies on its K4 residue, we profiled DNA methylation levels by bisulfite sequencing (BS-seq). At H3K4me3-enriched genes, gene body CG methylation levels were slightly reduced in *h3.3ko;HTR5* compared with WT (Supplemental Figure 4c), likely due to the absence of the other two H3.3 coding genes, *HTR4* and *HTR8*. Nevertheless, the K4A mutation in H3.3 did not further reduce CG methylation levels. Likewise, WT and the *sdg2* mutant showed comparable DNA methylation levels (Supplemental Figure 4c). Similarly, when non H3K4me3-enriched genes or transposable elements (TEs) were evaluated, DNA methylation levels in *h3.3ko;HTR5 K4A* and *sdg2* were similar to those of *h3.3ko;HTR5* and WT, respectively (Supplemental Figure 4c). Thus, the H3.3K4 residue and SDG2 are overall not necessary for DNA methylation in *Arabidopsis*.

We recently reported that a complete loss of H3.3 results in strong alterations in chromatin accessibility, with accessible levels at the gene 5’ regions being drastically reduced in *h3.3ko*, while the gene 3’ regions gain accessibility ^59^. To address whether the H3.3K4 residue plays a role in controlling chromatin accessibility, we analysed genome-wide chromatin accessibility by assay for transposase-accessible chromatin with sequencing (ATAC-Seq). Genes enriched with H3K4me3 were much more accessible than non H3K4me3-enriched genes (Supplemental Figure 4d), consistent with the notion that H3K4me3 is associated with active transcription. Due to the partial loss of H3.3 (*HTR4* and *HTR8*), *h3.3ko;HTR5* showed a mildly decrease and increase in chromatin accessibility at the gene 5’ and 3’ regions, respectively (Supplemental Figure 4d). The H3.3 K4A mutation further caused a slight reduction in chromatin accessibility at the gene 5’ regions, which also became less accessible in *sdg2* (Supplemental Figure 4d). These results suggest that H3K4me3 plays a minor role in chromatin accessibility control.

### H3.3K4 mutation induces similar transcriptome changes as in *sdg2*

To assess whether SDG2-mediated gene expression is regulated by H3.3K4, we analysed transcriptome changes in *h3.3ko;HTR5 K4A* and *sdg2* by RNA-seq. Given that transcriptome changes in *h3.3ko;HTR5 K4A* and *sdg2* are supposed to be measured by comparing with *h3.3ko;HTR5* and WT Col, respectively, hindering their direct comparison, we first compared the transcriptome of Col with *h3.3ko;HTR5*. Limited differences between them were detected (Figure 3a and 3b), consistent with their comparable phenotypes and H3K4 methylation levels. Subsequently, we compared the transcriptomes of *h3.3ko;HTR5 K4A* and *sdg2* both to WT. Overall, gene transcript level changes in *h3.3ko;HTR5 K4A* and *sdg2* compared with WT were highly correlated, with thousands of genes being significantly misexpressed (fold change >2 and *P*-adjust < 0.05) (Figure 3b and 3c). These misexpressed genes in *h3.3ko;HTR5 K4A* and *sdg2* showed strong overlap, and they were prevalently enriched in responsive, growth regulation and photosynthesis pathways (Figure 3d and Supplemental Figure 5a and 5b). Together, these results are in line with the strong reduction of H3K4me3 in both *h3.3ko;HTR5 K4A* and *sdg2*.

**Figure 3.**
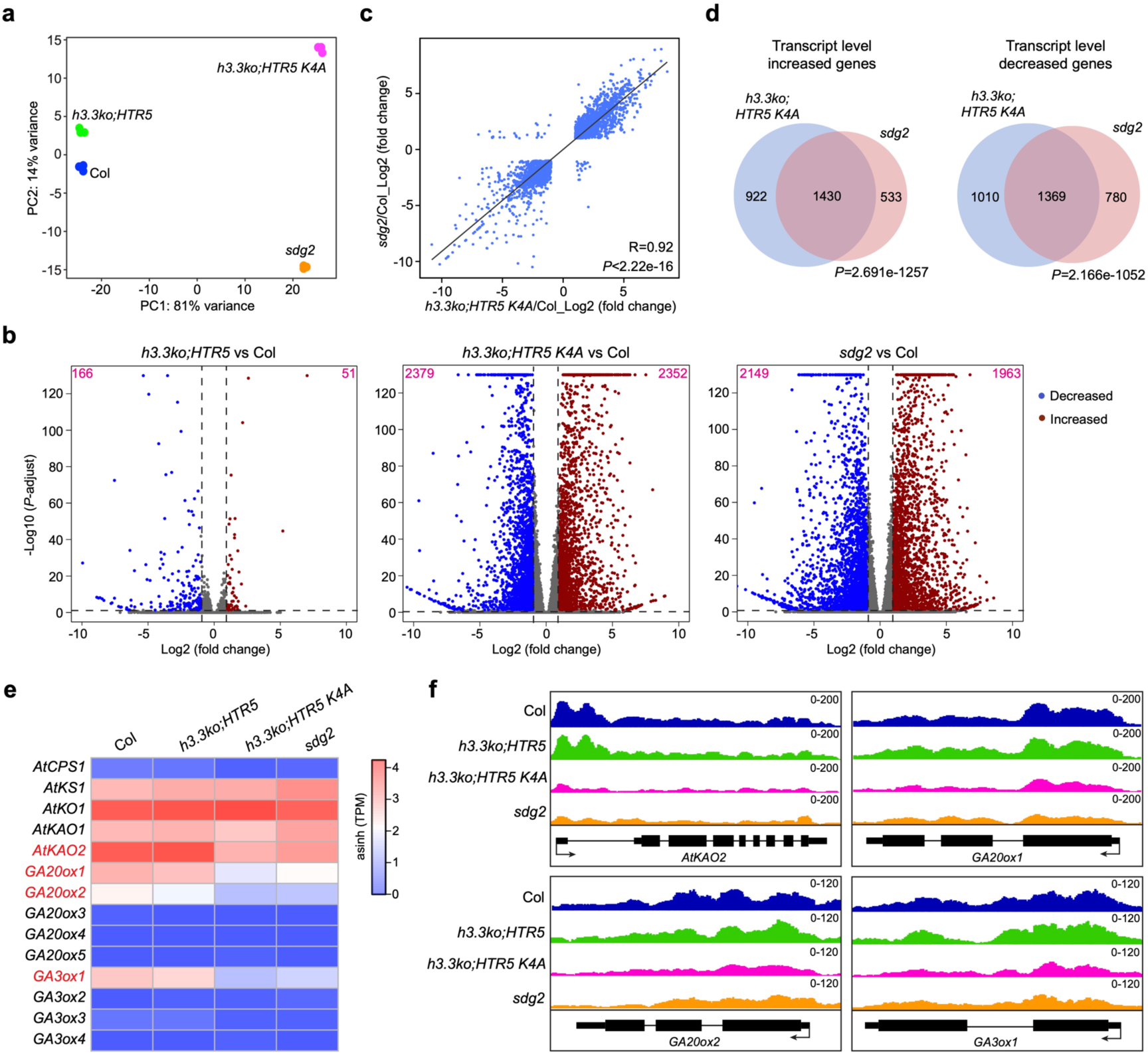
*h3.3ko;HTR5 K4A* shares transcriptional similarity with *sdg2*. **a.** PCA plot of Col, *h3.3ko;HTR5*, *h3.3ko;HTR5 K4A* and *sdg2* showing transcriptome differences determined by RNA-seq. PC1 covers the highest amount of variance between samples, PC2 covers most of the remaining variance. Three biological replicates were performed for each line. **b.** Volcano plots of differentially expressed genes (DEGs). The y-axis values correspond to -log_10_ (*P*-adjust), and the x-axis values correspond to log_2_ (fold change). Genes with at least two-fold expression changes and *P*-adjust less than 0.05 are considered misexpressed. The numbers of transcript level increased and decreased genes were indicated at the top right and left corners, respectively. **c.** Correlation of changes in transcript levels of significantly misexpressed genes in *h3.3ko;HTR5 K4A* and *sdg2* compared to Col. **d.** Venn diagrams of transcript level significantly increased and decreased genes in *h3.3ko;HTR5 K4A* and *sdg2* compared to Col. *P* values were calculated with hypergeometric test. **e.** Heatmap showing transcript levels of GA biosynthetic genes determined by RNA-seq. **f.** Genome browser view of H3K4me3 accumulation levels at *AtKAO2*, *GA20ox1*, *GA20ox2* and *GA3ox1* in Col, *h3.3ko;HTR5*, *h3.3ko;HTR5 K4A* and *sdg2*.

**Figure 4.**
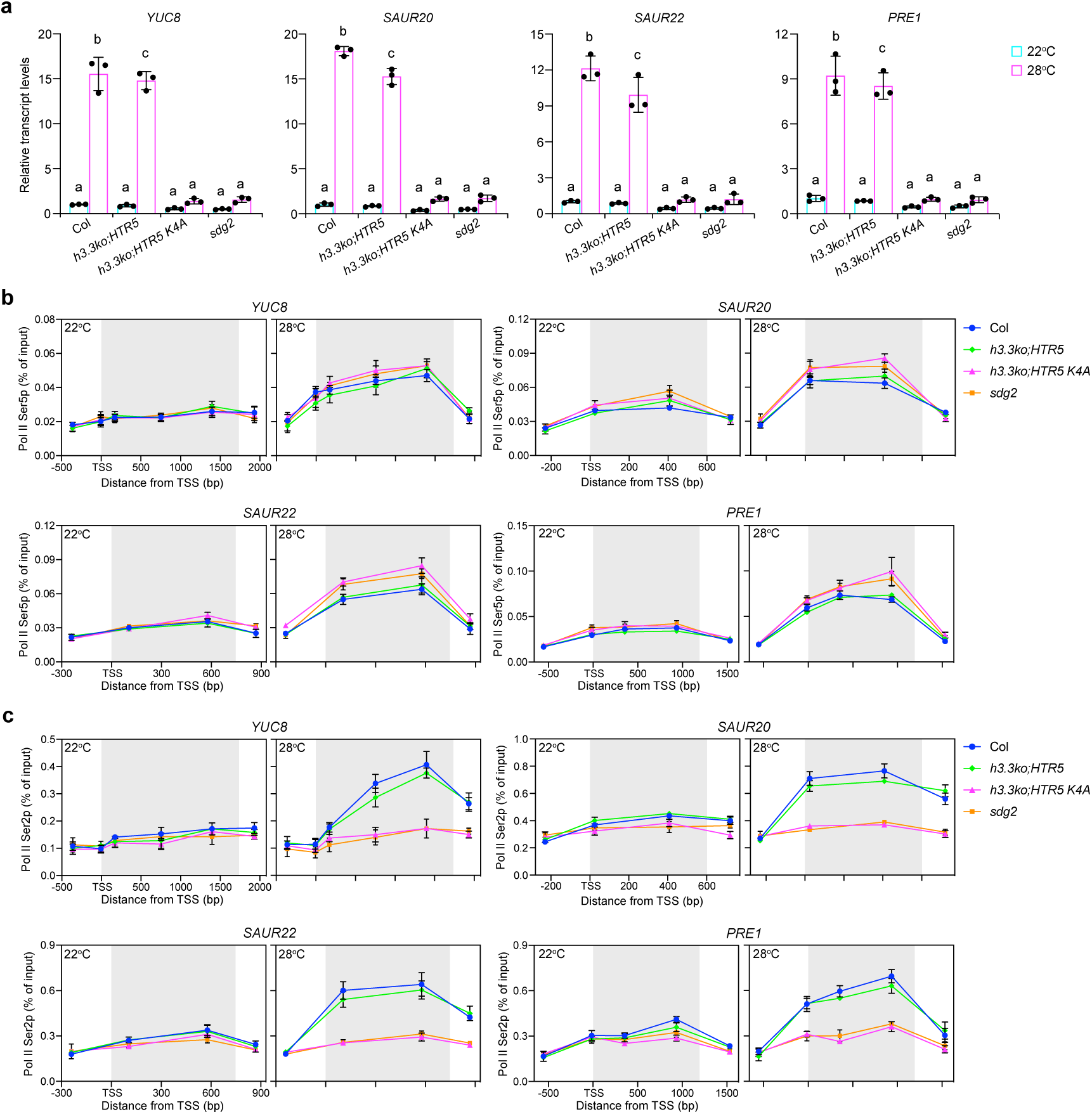
Loss of H3.3K4 or SDG2 impairs high temperature-induced de novo gene activation and RNA Pol II elongation. **a.** Relative transcript levels of *YUC8*, *SAUR20*, *SAUR22* and *PRE1* determined by RT-qPCR at 22°C and 28°C. Transcript levels in Col, 22°C were set as 1. Values are means ± SD of three biological replicates. The significance of differences was tested using one-way ANOVA with Tukey’s test (*P* < 0.05), with different letters indicating statistically significant differences. **b and c.** RNA Pol II Ser5p (b) and RNA Pol II Ser2p (c) enrichment levels across *YUC8*, *SAUR20*, *SAUR22* and *PRE1* determined by ChIP-qPCR at 22°C and 28°C. Values are means ± SD of three biological replicates. Genic regions are indicated with grey shading.

**Figure 5.**
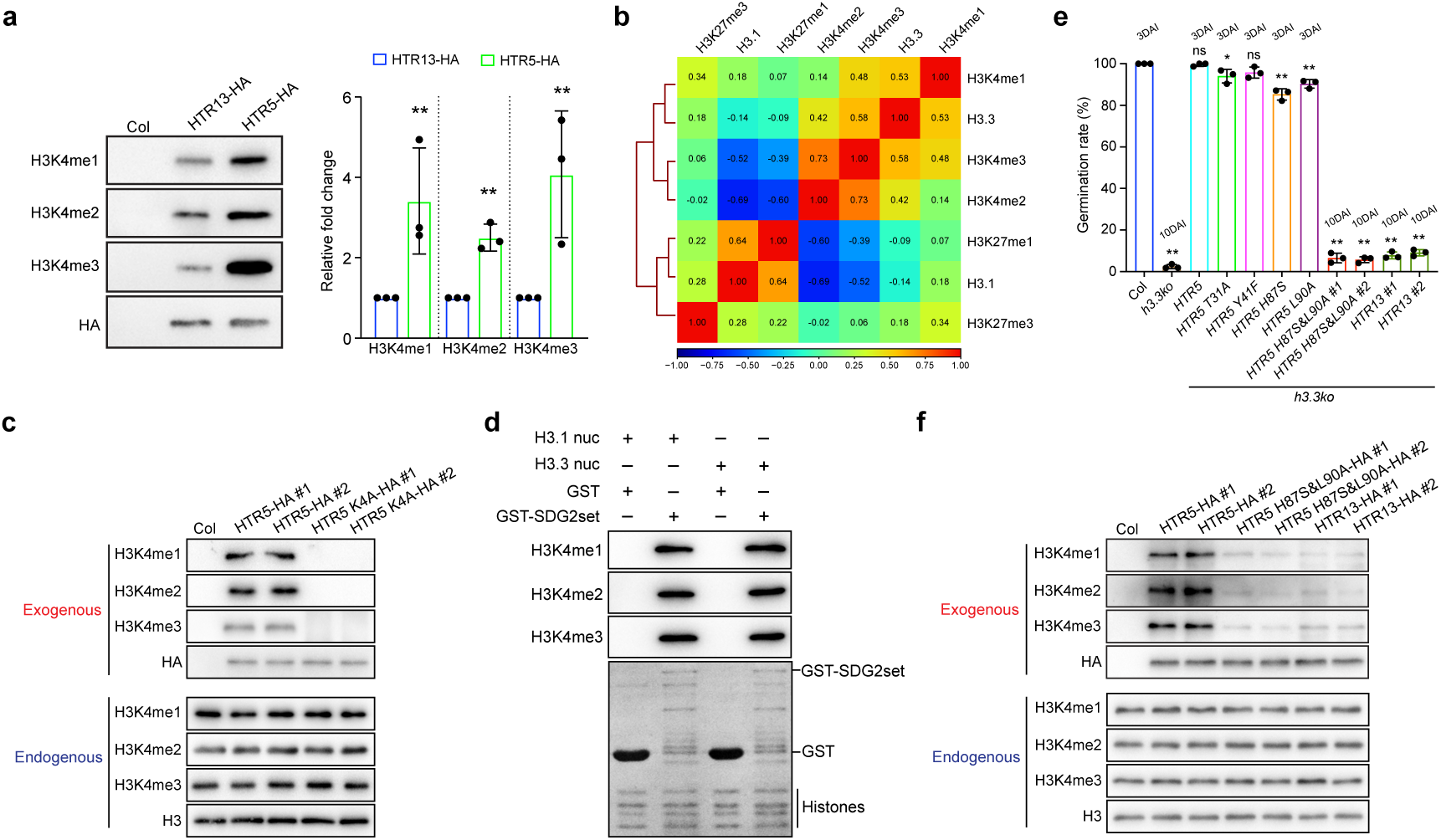
H3K4 methylation is preferentially enriched on H3.3 over H3.1. **a.** H3K4 methylation levels on exogenous H3.1 (HTR13) and H3.3 (HTR5) determined by western blotting. The bar chart represents quantification of western blot signals from three biological replicates (Figure 5a and Supplemental Figure 7a). Values are means ± SD. The significance of differences was determined by two-tailed Student’s *t*-test (**, *P* < 0.01). **b.** Heatmap showing the Pearson correlation between genome-wide distributions of indicated histone variants and modifications profiled by ChIP-seq. **c.** H3K4 methylation levels on exogenous H3.3 (HTR5) and H3.3 K4A (HTR5 K4A), and endogenous H3 determined by western blotting. Results from two independent transgenic lines are shown. **d.** *In vitro* methyltransferase assay showing the activity of SDG2 SET domain (SDG2set) on nucleosomes containing H3.1 or H3.3. The levels of H3K4 methylation in *in vitro* methyltransferase assay products were determined by western blotting. **e.** Seed germination rates of indicated lines after imbibed for 3 days (3DAI) or 10 days (10DAI). Values are means ± SD of three biological replicates. At least 53 seeds were analysed per replicate. Statistical significance relative to Col was determined by two-tailed Student’s *t*-test (**, *P* < 0.01; ns, not significant). **f.** H3K4 methylation levels on exogenous H3.3 (HTR5), H3.3 H87S&L90A (HTR5 H87S&L90A), H3.1 (HTR13), and endogenous H3 determined by western blotting. Results from two independent transgenic lines are shown.

Both *h3.3ko;HTR5 K4A* and *sdg2* exhibited reduced growth and dwarfism. Gibberellic acid (GA) is a crucial plant hormone that stimulates growth and development ^74,75^. Loss of GA biosynthetic enzymes, such as GA20ox1, GA20ox2, or GA3ox1, results in dwarf phenotypes ^76,77^. We thus examined the transcript levels of GA biosynthetic genes ^78^. Several of them, including *AtKAO2*, *GA20ox1*, *GA20ox2*, and *GA3ox1*, showed reduced transcript levels in *h3.3ko;HTR5 K4A* and *sdg2* compared with WT and *h3.3ko;HTR5* (Figure 3e), providing a potential explanation for the observed growth phenotypes. Moreover, in agreement with their reduced transcription, H3K4me3 enrichment levels at their loci were diminished in *h3.3ko;HTR5 K4A* and *sdg2* (Figure 3f).

H3K4me3 has been implicated in pre-mRNA splicing ^79,80^. In *Arabidopsis*, the COMPASS-like complex interacts with the mRNA cap binding complex to regulate intron splicing ^81^. By analysing the RNA-seq data, we found that a number of introns showed significant intron retention (IR) changes in *h3.3ko;HTR5 K4A* and *sdg2* (Supplemental Figure 5c), with a notable overlap between them. (Supplemental Figure 5d). Most IR changes in *h3.3ko;HTR5 K4A* and *sdg2* occurred at 5’ proximal introns (Supplemental Figure 5e), consistent with the enrichment of H3K4me3 around the TSS. Taken together, our results show that *h3.3ko;HTR5 K4A* and *sdg2* share similar gene expression programs.

### H3.3K4 and SDG2 are required for de novo gene activation and transcriptional elongation

It is noteworthy that although H3K4me3 is a permissive histone modification, the number of significantly downregulated genes in *h3.3ko;HTR5 K4A* and *sdg2* is not substantially higher than the number of upregulated genes (Figure 3b). Moreover, the reduction in H3K4me3 levels was only mildly correlated with changes in gene expression (Supplemental Figure 6a and 6b). We further conducted ChIP-seq to profile RNA Pol II CTD Ser5 phosphorylation (Pol II Ser5p), the initiation and pausing form, and the RNA Pol II CTD Ser2 phosphorylation (Pol II Ser2p), the elongation form. However, their overall enrichment at H3K4me3-marked genes also showed no strong changes in *h3.3ko;HTR5 K4A* and *sdg2* (Supplemental Figure 6c and 6d).

**Figure 6.**
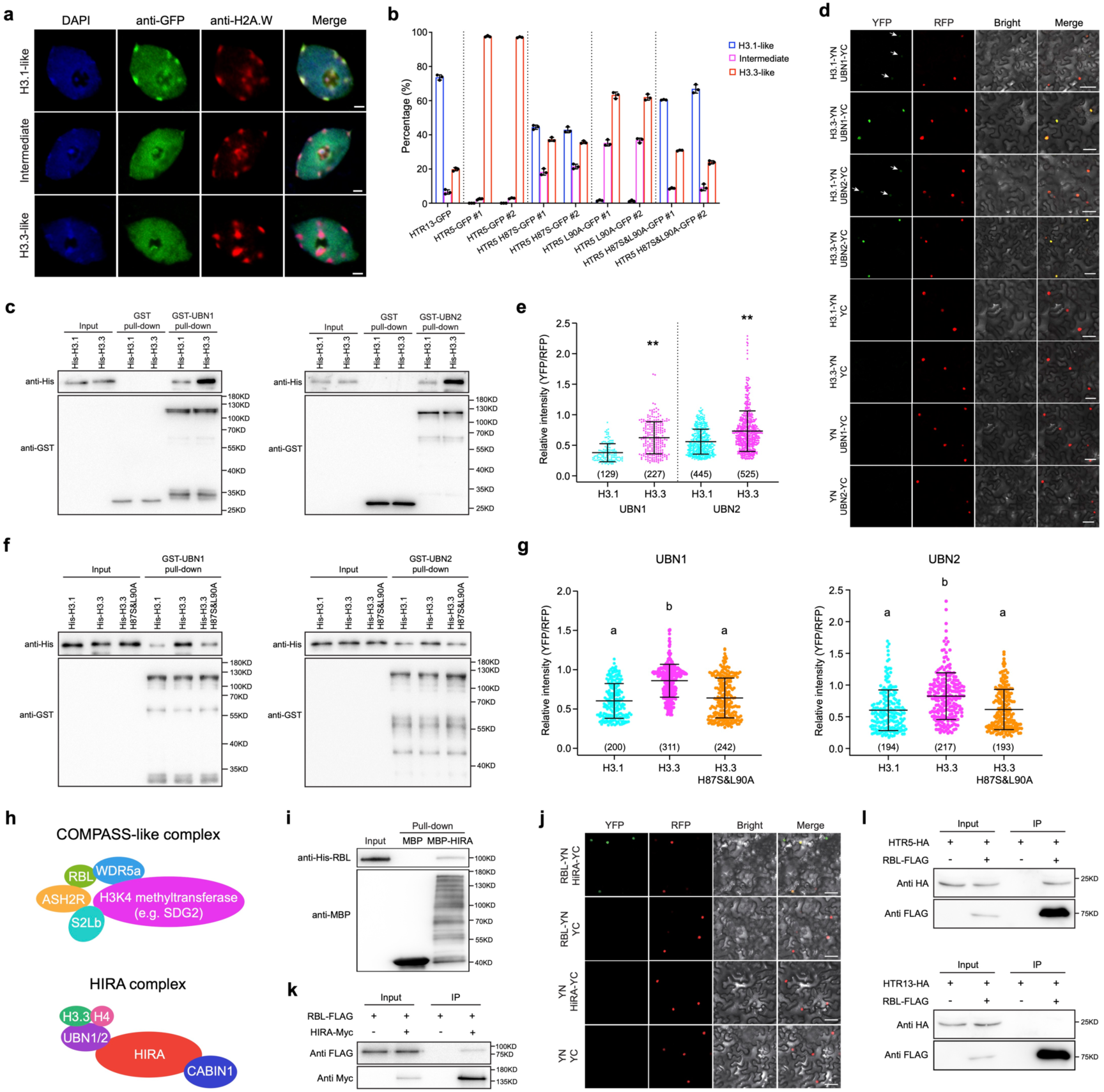
H3K4 methylation is closely linked with the deposition of H3.3. **a.** Distribution patterns of GFP-fused H3.1 (HTR13) or H3.3 (HTR5) in leaf nuclei. Condensed chromocenters were immunostained with the heterochromatin mark H2A.W. Scale bars, 2 μm. **b.** Percentages of nuclei showing H3.1-like, intermediate and H3.3-like distribution patterns for the indicated proteins. Results from two independent transgenic lines are shown. Values are means ± SD of three biological replicates. At least 91 nuclei were analysed per replicate. **c.** Pull-down assay of GST-UBN1/2 with His-H3.1 or His-H3.3. GST or GST-UBN1/2 was incubated with an equal amount of His-H3.1 or His-H3.3. Proteins were recovered with glutathione-agarose resin and analysed by immunoblotting with anti-His or anti-GST antibody. **d.** BiFC assay showing representative fluorescence signals of UBN1/2-cYFP (YC) with H3.1-nYFP (YN) or H3.3-YN in *N. benthamiana* leaf cells. Arrows indicate nuclei with the YFP fluorescence signals for UBN1/2-YC with H3.1-YN. A nucleus localized protein mRFP-AHL22 was employed as a control. Same amount of agrobacterium containing YN, YC and mRFP-AHL22 were used in each transformation, and YFP and RFP signals were acquired using a Zeiss confocal laser-scanning microscope with the same setting across all transformations. Scale bars, 50 μm. **e and g.** Relative intensity of YFP signals compared to RFP (mRFP-AHL22) as a control in the BiFC assay. Numbers in parentheses indicate the amounts of quantified nuclei. The significance of differences was determined by two-tailed Student’s *t*-test (**, *P* < 0.01) (e) or by one-way ANOVA with Tukey’s test (*P* < 0.05), with different letters indicating statistically significant differences (g). **f.** Pull-down assay of GST-UBN1/2 with His-H3.1, His-H3.3 or His-H3.3 H87S&L90A. GST or GST-UBN1/2 was incubated with an equal amount of His-H3.1, His-H3.3 or His-H3.3 H87S&L90A. Proteins were recovered using glutathione-agarose resin and analysed by immunoblotting with anti-His or anti-GST antibody. **h.** Schematic diagram of *Arabidopsis* COMPASS-like and HIRA complex components. **i.** Pull-down assay of MBP-HIRA with His-RBL. MBP or MBP-HIRA was incubated with His-RBL. Proteins were recovered with amylose resin and analysed by immunoblotting with anti-His or anti-MBP antibody. **j.** BiFC assay showing the interaction of RBL-YN with HIRA-YC in *N. benthamiana* leaf cells. A nucleus localized protein mRFP-AHL22 was used for nuclei labeling. Scale bars, 50 μm. **k.** Co-IP assay of RBL with HIRA. Total proteins were extracted from *Arabidopsis* plants expressing both RBL-FLAG and HIRA-Myc, or only RBL-FLAG. After recovered with anti-Myc beads, proteins were analysed with anti-FLAG or anti-Myc antibody. **l.** Co-IP assay of RBL with H3.1 or H3.3. Total proteins were extracted from *Arabidopsis* plants expressing RBL-FLAG with either H3.1 (HTR13)-HA, H3.3 (HTR5)-HA, or only H3.1 (HTR13)-HA or H3.3 (HTR5)-HA. After recovered with anti-FLAG beads, proteins were analysed with anti-FLAG or anti-HA antibody.

Transcription changes upon the removal of H3K4me3 under steady-state may not fully reflect the direct role of H3K4me3 in gene expression regulation ^82,83^. Indeed, recent studies inducing a rapid depletion of COMPASS-like complex subunits in mouse embryonic stem cells have demonstrated that most genes misexpressed in response to acute loss of H3K4me3 are downregulated, supporting the role of H3K4me3 in promoting transcription ^83,84^. We thus tested whether H3K4me3 is necessary for de novo gene activation induced by high temperatures. We decided to perform high temperature treatment for several reasons. Firstly, elevated temperatures activate many auxin-related genes (e.g. *YUC8*, *SAUR20*, *SAUR22* and *PRE1*), which are important for plant adaptation to high temperatures ^85^. Secondly, this gene activation is associated with the deposition of H3.3 and H3K4me3 at their loci ^57,86^. Lastly, H3.3 has been shown to be required for gene activation under high temperature conditions ^57^. Upon high temperature treatment, the transcription of *YUC8*, *SAUR20*, *SAUR22*, and *PRE1* was strongly activated in Col and *h3.3ko;HTR5*. However, their transcript levels were much lower in *h3.3ko;HTR5 K4A* and *sdg2* (Figure 4a). This aligns with the absence of high temperature-induced H3K4me3 deposition at their loci in *h3.3ko;HTR5 K4A* and *sdg2* (Supplemental Figure 6e and 6f). In addition, we observed an accumulation of Pol II Ser5p, while levels of Pol II Ser2p did not increase with high temperature treatment (Figure 4b and 4c). These results highlight the crucial role of H3K4me3 in facilitating de novo gene activation and RNA Pol II elongation.

### H3K4 methylation is preferentially enriched on H3.3

Both H3.1 and H3.3 bear the K4 residue, but K4 is essential only for the function of H3.3, not H3.1 (Figure 1 and Supplemental Figure 1 and 2). Furthermore, loss of H3.3K4 strongly diminishes global H3K4 methylation levels (Figure 2). To explore the underlying mechanism, we employed transgenic lines expressing HA-tagged H3.1 (HTR13) or H3.3 (HTR5) with their respective promoters in WT, so that the exogenous H3.1/H3.3 can be distinguished from endogenous H3 based on protein size differences. Notably, H3.3 exhibited higher levels of all three forms of H3K4 methylation compared to H3.1 (Figure 5a and Supplemental Figure 7a). Consistent with this, H3.3 is more enriched in regions marked by H3K4 methylation, while the genomic distribution of H3.1 preferentially correlates with repressive modifications like H3K27me1 and H3K27me3 (Figure 5b, and Supplemental Figure 7b-7d) ^38^. We further introduced HA tag-fused H3.3 or H3.3 K4A with the *HTR5* promoter into WT and examined H3K4 methylation levels on both endogenous H3 and exogenous H3.3. Introducing K4A mutated H3.3 did not affect the endogenous H3K4 methylation levels, indicating that mutating K4 to A in H3.3 does not interfere with H3K4 methylation in trans (Figure 5c). This is in agreement with the observations that H3.3 K4A did not induce gain-of-function effects (Supplemental Figure 2f). Hence, our data reveal that H3.3 serves as an important carrier of H3K4 methylation.

**Figure 7.**
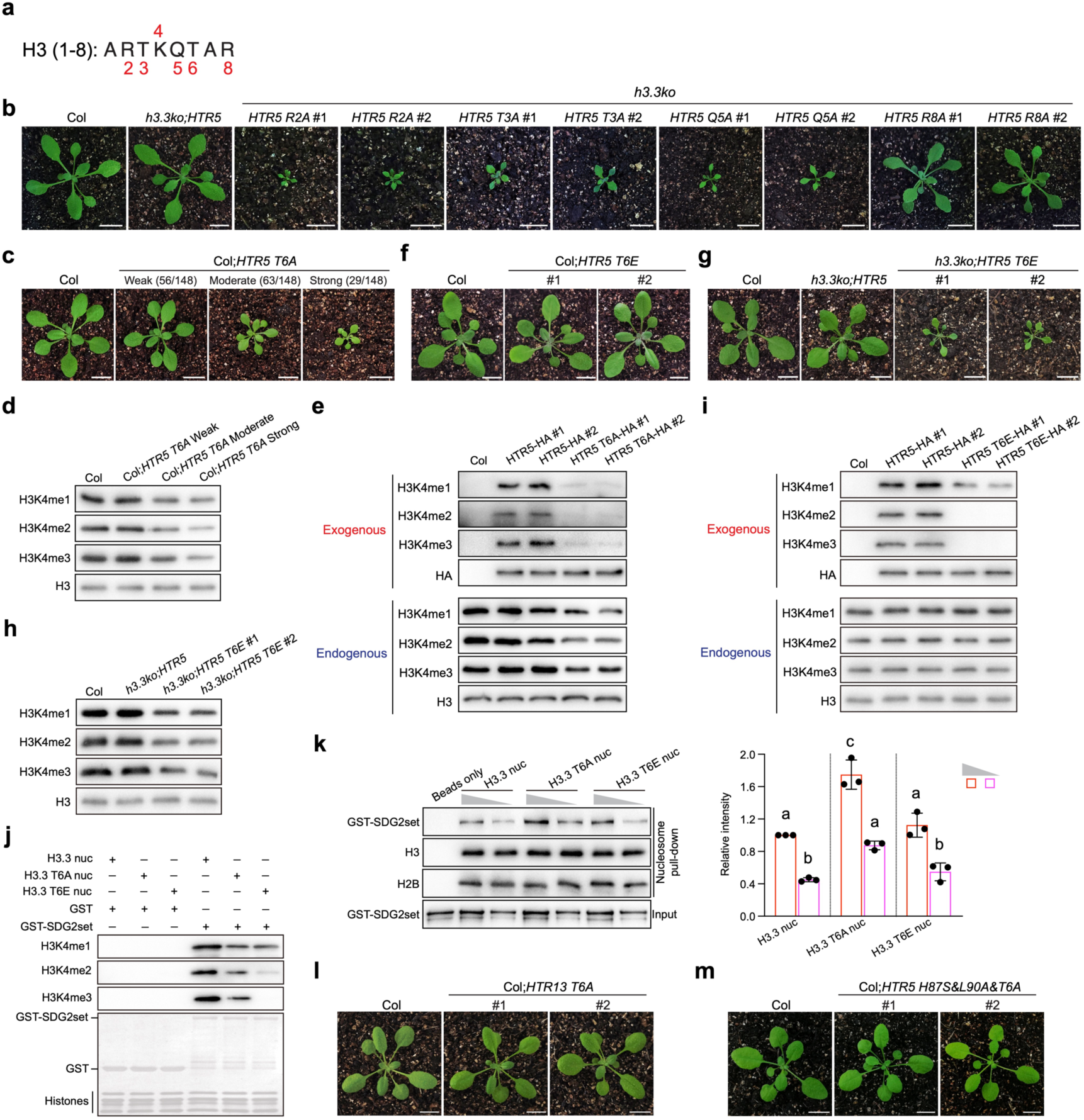
Mutations at H3.3K4 neighbouring residues affect H3K4 methylation and the function of H3.3. **a.** Diagram of H3.3K4 neighbouring non-alanine amino acids that were mutated in this study. **b.** Developmental phenotypes of the indicated lines at the vegetative stage. Two independent representative transgenic lines are shown. Scale bars, 1 cm. **c.** Developmental phenotypes of T1 transgenic lines expressing *HTR5 T6A* in the Col background at the vegetative stage. Representative plants showing weak, moderate and strong phenotypes are shown. A total of 148 T1 plants were generated, with the numbers of plants exhibiting weak, moderate and strong phenotypes indicated in parentheses. **d.** H3K4 methylation levels in the indicated T1 transgenic lines determined by western blotting. H3 was employed as a loading control. **e.** H3K4 methylation levels on exogenous H3.3 (HTR5), H3.3 T6A (HTR5 T6A), and endogenous H3 determined by western blotting. Results from two independent transgenic lines are shown. **f and g.** Developmental phenotypes of transgenic lines expressing *HTR5 T6E* in the Col (f) or *h3.3ko* (g) background at the vegetative stage. Two independent representative transgenic lines are shown. Scale bars, 1 cm. **h.** H3K4 methylation levels in the indicated lines determined by western blotting. H3 was employed as a loading control. Results from two independent transgenic lines are shown. **i.** H3K4 methylation levels on exogenous H3.3 (HTR5), H3.3 T6E (HTR5 T6E), and endogenous H3 determined by western blotting. Results from two independent transgenic lines are shown. **j.** *In vitro* methyltransferase assay showing the activity of SDG2 SET domain (SDG2set) on nucleosomes containing H3.3, H3.3 T6A or H3.3 T6E. The levels of H3K4 methylation in *in vitro* methyltransferase assay products were determined by western blotting. **k.** Mononucleosome pull-down assay of H3.3, H3.3 T6A and H3.3 T6E containing nucleosome incubated with GST-SDG2 SET domain (SDG2set) proteins. H3 and H2B were employed as loading controls. The bar chart represents quantification of western blot signals from three independent replicates (Figure 6k and Supplemental Figure 13i). Values are means ± SD, normalized with H3 signals. The significance of differences was determined by one-way ANOVA with Tukey’s test (*P* < 0.05), with different letters indicating statistically significant differences. **l and m.** Developmental phenotypes of transgenic lines expressing *HTR13 T6A* (l) or *HTR5 H87S&L90A&T6A* (m) in the Col background at the vegetative stage. Two independent representative transgenic lines are shown. Scale bars, 1 cm.

To investigate why H3K4 methylation is preferentially enriched on H3.3 over H3.1, we performed an *in vitro* methylation assay using the catalytic SET domain of SDG2 and reconstituted nucleosomes containing either H3.1 or H3.3. We also included the SET domain of ATXR6, a plant-specific H3K27 methyltransferase selectively catalyses H3K27me1 on H3.1 as a control ^61^. Our experiments successfully replicated the specific activity of ATXR6 (Supplemental Figure 7e). However, SDG2 did not exhibit preferential methylation on H3.1 or H3.3 (Figure 5d). These results suggest that SDG2 does not selectively methylate H3.1 or H3.3 *in vitro*, though we could not exclude the possibility that other H3K4 methyltransferases might preferentially methylate H3.3 over H3.1, or that H3.1 could be a more favoured substrate for H3K4 demethylation compared to H3.3.

### H87 and L90 residues in H3.3 are crucial for its function and H3K4 methylation

We hypothesized that the four amino acids differing between H3.3 and H3.1 may contribute to their differential H3K4 methylation levels (Figure 1a). To test this possibility, we mutated each of these four amino acids in H3.3 (HTR5) to their corresponding amino acids in H3.1, and performed complementation tests by expressing these mutated H3.3 with the *HTR5* promoter in *h3.3ko*. None of the single amino acid mutations strongly affected the function of H3.3 in germination and plant development (Figure 5e and Supplemental Figure 8a), yet H3.3 with the H87S or L90A mutation showed slightly lower ability to rescue the germination defects of *h3.3ko* compared to H3.3 with T31A or Y41F mutation (Figure 5e). We therefore mutated both H87 and L90 in H3.3, and in this case the function of mutated H3.3 in germination control is severely impaired (Figure 5e). Expressing H3.1 (HTR13) with the *HTR5* promoter in *h3.3ko* also resulted in similar germination defects (Figure 5e) ^59^. Together, we conclude that the H87 and L90 amino acids are crucial for the function of H3.3.

To assess the involvement of these two residues in H3K4 methylation, HA tag-fused H3.3, H3.3 H87S&L90A, or H3.1 was expressed with the *HTR5* promoter in WT. Compared to H3.3, both H3.1 and the H87 and L90 mutated H3.3 carried lower levels of H3K4 methylation, but they did not affect endogenous H3K4 methylation (Figure 5f). This suggests that the H87 and L90 amino acids in H3.3 determine the enrichment of H3K4 methylation in cis.

### H87 and L90 residues in H3.3 control its chromatin deposition that is linked with H3K4 methylation

Since H87 and L90 residues are located within the core of H3.3 and are distant from K4, they are unlikely to directly influence H3K4 methylation or demethylation activity. In animals, H3.3-specific residues A87, I89 and G90 regulates its deposition ^43^. Moreover, mutating H87 and L90 residues in *Arabidopsis* H3.3 to their H3.1 counterparts led to an H3.1-like localization pattern at the rDNA loci when the mutated H3.3 was transiently expressed in tobacco cells ^71^. To assess the importance of H87 and L90A in the chromatin deposition of H3.3 in *Arabidopsis*, we fused H3.3, H3.3 H87S, H3.3 L90A, or H3.3 H87S&L90A with GFP, and expressed them in WT with the *HTR5* promoter. In all cases, GFP signals were detected within nuclei (Supplemental Figure 8b), showing that H87 and L90 are not required for the subcellular localization of H3.3. We then performed immunostaining to examine their chromatin localization. In general, H3.1 is distributed across the genome with stronger enrichment at the heterochromatic regions, while H3.3 is primarily localized to euchromatic regions ^38,39,71,87^. In an H3.1 (HTR13)-GFP transgenic line ^39,87^, GFP signals in most nuclei showed typical H3.1-like distribution patterns (strong signals at the heterochromatin-enriched chromocenters), with a small portion of nuclei displayed H3.3-like patterns (no signal at the chromocenters) or an intermediated state (weak signals at the chromocenters) (Figure 6a and 6b). In H3.3-GFP transgenic lines, GFP signals exhibited almost exclusively H3.3-like distribution patterns (Figure 6a and 6b). Mutating H87 or L90 in H3.3 altered its chromatin distribution, with H3.3 bearing both mutations exhibiting patterns similar to H3.1 (Figure 6a and 6b). To confirm these findings, we further performed immunostaining analysis with transgenic lines expressing HA-tagged H3.1, H3.3, or H3.3 H87S&L90A (Figure 5f), and observed similar results (Supplemental Figure 8c). Together, these results suggest that the H87 and L90 residues in H3.3 are critical for its chromatin distribution.

In animals, the A87 and G90 residues in H3.3 mediate its preferential binding with UBN1/2 and DAXX, which confer specificity for H3.3 deposition by the HIRA complex and ATRX-DAXX, respectively ^50,51,88,89^. However, DAXX is not conserved in plants ^45,47^. To test whether the H87 and L90 residues in *Arabidopsis* H3.3 have similar functions, we first examined the interaction of *Arabidopsis* UBN1/2 with H3.1 and H3.3. Pull-down assays showed that UBN1/2 preferentially binds to H3.3 over H3.1 (Figure 6c). A further test in planta using Bimolecular Fluorescence Complementation (BiFC) assay confirmed the selective association of UBN1/2 with H3.3 (Figure 6d and 6e). We then examined the binding of UBN1/2 to H3.3 with H87S and L90A mutations, and found that these mutations reduced their interactions (Figure 6f and 6g). These data suggest that the H87 and L90 residues in plant H3.3 are essential for its preferred binding with UBN1/2.

The above results suggest that the H87 and L90 residues in H3.3 are required for both its chromatin deposition and the enrichment of H3K4 methylation. Since H3.3 is not selectively methylated by SDG2, it is likely that its specific chromatin deposition contributes to the enrichment of H3K4 methylation. H3K4 methylation is catalysed by the COMPASS or COMPASS-like complex, which consists of an H3K4 methyltransferase (e.g. SDG2) and several other core components ^16,18,20,30^. We tested the interaction between the *Arabidopsis* COMPASS-like complex components and the HIRA complex subunits, and found that RBL directly interacted with HIRA in the pull-down assay (Figure 6h and 6i). This interaction was further confirmed by BiFC and co-immunoprecipitation (Co-IP) experiments (Figure 6j and 6k). In addition, a Co-IP assay of RBL-FLAG with H3.1 (HTR13)-HA or H3.3 (HTR5)-HA showed that RBL preferentially associated with H3.3 compared to H3.1 (Figure 6l). Collectively, these results provide evidence for a close link between the deposition of H3.3 and the catalysation of H3K4 methylation.

### Residues nearby H3.3K4 are essential for the H3.3 function

Given that K4 in H3.3 is essential for its function, we investigated the importance of K4 nearby non-alanine residues in H3.3, which could be subjected to modifications (e.g. methylation, serotonylation, or phosphorylation) or have been reported to regulate H3K4me3 levels or its recognition by proteins such as TFIID (Figure 7a) ^90-93^. Each of these residues was mutated to alanine and the mutated H3.3 was employed for the *h3.3ko* complementation. Plants complemented with *HTR5 R2A*, *HTR5 T3A*, or *HTR5 Q5A* showed strong developmental defects similar to those observed in *h3.3ko;HTR5 K4A* (Figure 7b and Supplemental Figure 9a-9c)*. h3.3ko;HTR5 R8A* also displayed moderate defects at the vegetative stage (Figure 7b and Supplemental Figure 9d). In addition, all these mutations caused early flowering and sterility (Supplemental Figure 9e and 9f). However, we were unable to obtain any complementation plants expressing *HTR5 T6A* (see below). In contrast, mutating these residues in H3.1 did not affect its function (Supplemental Figure 10), indicating that, like K4, these residues are specifically required for the function of H3.3.

To determine whether residues nearby K4 (excluding T6) regulate H3K4 methylation, we measured global H3K4 methylation levels in these complementation plants, and found that they were strongly reduced in *h3.3ko;HTR5 R2A* and *h3.3ko;HTR5 T3A* but not in *h3.3ko;HTR5 Q5A* and *h3.3ko;HTR5 R8A* (Supplemental Figure 11a and 11b). Expressing HA-tagged H3.3 R2A and H3.3 T3A in WT did not reduce endogenous H3K4 methylation levels (Supplemental Figure 11c), similar to what was observed with H3.3 K4A (Figure 5c). However, it is of note that the R2A or T3A mutation partially affects the affinity of the anti-H3K4me3 antibody (Supplemental Figure 11d). Therefore, the significance of H3.3R2 and H3.3R2T3 in H3K4me3 accumulation remains uncertain, and they may also influence the reading of H3K4me3 ^94^. The R2 and T3 residues can be methylated and phosphorylated, respectively ^95,96^, and the latter could be mimicked by glutamic acid (E) ^97^. We found that *h3.3ko;HTR5 T3E* plants showed similar developmental defects as *h3.3ko,HTR5 T3A* (Supplemental Figure 12). This observation is in agreement with the notion that phosphorylation at H3T3 anticorrelates with H3K4me3 and decreases its association with TFIID ^93,96,98^. Hence, it seems that only the unphosphorylated T3 (without being mutated to A or phosphorylated) is permissive for H3K4me3 deposition and/or its recognition.

### The H3.3 T6A mutation causes gain-of-function effects that impair SDG2 activity

Because we were unable to obtain *h3.3ko* plants expressing *HTR5 T6A*, we examined the phenotypes of *h3.3ko/+*;*HTR5 T6A*. Unlike other mutations, the T6A mutated H3.3 caused severe developmental defects including dwarfism and sterility, even in *h3.3ko/+* (Supplemental Figure 13a-13c). This suggests that T6A mutation in H3.3 may induce gain-of-function effects. To test this possibility, we expressed *HTR5 T6A* in the WT Col background with the *HTR5* promoter. T1 transgenic plants showed a range of weak to severe developmental defects, with the more severe cases being completely sterile (Figure 7c and Supplemental Figure 13d). Moreover, global H3K4 methylation levels were reduced, particularly in plants with moderate and strong defects (Figure 7d). To investigate how the H3.3 T6A mutation affects H3K4 methylation, we expressed HA-tagged H3.3 T6A in WT, and found that both the exogenous H3.3 and endogenous H3 lost H3K4 methylation (Figure 7e). These results indicate that the T6A mutation in H3.3 also impairs H3K4 methylation in trans.

We then mutated the H3.3 T6 residue to E, which can mimic phosphorylation. Expressing *HTR5 T6E* in WT did not induce developmental abnormities (transgenic plants n=56) (Figure 7f and Supplemental Figure 13e). However, H3.3 T6E was still not fully capable to rescue the *h3.3ko* (Figure 7g and Supplemental Figure 13f-13h), and *h3.3ko;HTR5 T6E* exhibited reduced global H3K4 methylation levels (Figure 7h). Unlike the T6A mutation, expressing H3.3 T6E did not notably affect H3K4 methylation levels on endogenous H3 (Figure 7i), suggesting that the T6E mutation only affects H3K4 methylation in cis. *In vitro* methylation assay revealed that both the T6A and T6E mutations impair SDG2 activity, with T6E showing a stronger inhibitory effect, particularly on H3K4me2 and H3K4me3 (Figure 7j).

Despite both mutations affecting SDG2 activity, only the T6A mutation in H3.3 induced a dominant effect. A gain-of-function K to M mutation at H3K27 is linked with certain pediatric brain cancers and significantly impairs the activity of polycomb repressive complex 2 (PRC2) that deposits H3K27 methylation ^65,99^. It is suggested that K27M retains and sequesters PRC2, as PRC2 shows a higher affinity for K27M compared with K27 ^99,100^. To test whether the T6A mutation exhibits a similar effect, we performed a mononucleosome pull-down assay and found that SDG2 indeed has a higher affinity for nucleosome containing H3.3 T6A compared to H3.3 and H3.3 T6E (Figure 7k and Supplemental Figure 13i). Structural predications also suggest that A6 residue might form additional hydrogen bond interaction with the amino acid in SDG2 SET domain (Supplemental Figure 14a-14c). These results hint a possibility that the T6A mutated H3.3 may retain and compromise SDG2 upon interaction, leading to the loss of H3K4 methylation in a dominant-negative fashion. Notably, expressing T6A mutated H3.1 (transgenic plants n=67) or T6A mutated H3.3 H87S&L90A (transgenic plants n=65) with the *HTR5* promoter in WT did not cause developmental defects (Figure 7l and 7m and Supplemental Figure 14d and 14e), further supporting the selective association between H3.3 and the H3K4 methylation machinery.

## Discussion

Histone modifications and variants greatly expand the chromatin diversity, providing broad mechanisms for chromatin regulation. However, their functional connections remain largely elusive. The incorporation of histone variants may directly impact on the chromatin activity owing to their sequence variations that may alter nucleosome properties. In addition, studies, primarily in animals, have suggested that various histone modifications show differential enrichment on histone variants ^101,102^. Several models have been proposed to explain this specificity. Some variants may possess unique residues that can be modified, such as S31 in animal H3.3 and the SQ motif in H2A.X ^103,104^. In other cases, sequence differences proximate to the modified residue may enhance or block the enzymatic activity despite the presence of the modified residue ^61,105^. In addition, nearby modifications could be prerequisites for other modifications. For instance, phosphorylation at S31 of animal H3.3 can stimulate acetylation at its K27 or trimethylation at its K36 ^55,60^.

In this work, we demonstrate that the K4 residue of H3.3 is preferentially methylated compared to H3.1 in *Arabidopsis* (Figure 5a). Consistently, K4 is essential for the function of H3.3 but not H3.1 (Figure 1 and Supplemental Figure 1 and 2). Interestingly, SDG2, a major H3K4 methyltransferase in *Arabidopsis*, does not directly differentiate H3.3 and H3.1, as suggested by its equal *in vitro* methylation activity on both (Figure 5d). Instead, the H3.3-specific residues H87 and L90, which determine the preferential binding of UBN1/2 to H3.3 (Figure 6c-6g, 8a), contribute to the predominant accumulation of H3K4 methylation on H3.3 (Figure 5f). Switching the H3.3 H87 and L90 residues to their counterparts in H3.1 leads to an H3.1-like distribution pattern and severely impairs H3.3 function (Figure 5e, 6a, 6b and Supplemental Figure 8c), supporting the idea that H87 and L90 in H3.3 determine its specific deposition by the HIRA complex. Considering that HIRA directly associates with RBL, a core subunit of the COMPASS-like complex (Figure 6h-6k), it is likely that H3K4 methylation is closely linked with H3.3 deposition, thereby enriching H3K4 methylation on H3.3 (Figure 8b). Notably, the amino acids at same positions in animal H3.3 (A87 and G90) are also required for its specific deposition and recognition by UBN1/2 ^43,50,51^. These findings may partially explain the convergent evolution of H3.1 and H3.3 in plants and animals, which, despite evolving independently in separate kingdoms, have acquired similar features ^106-108^.

**Figure 8.**
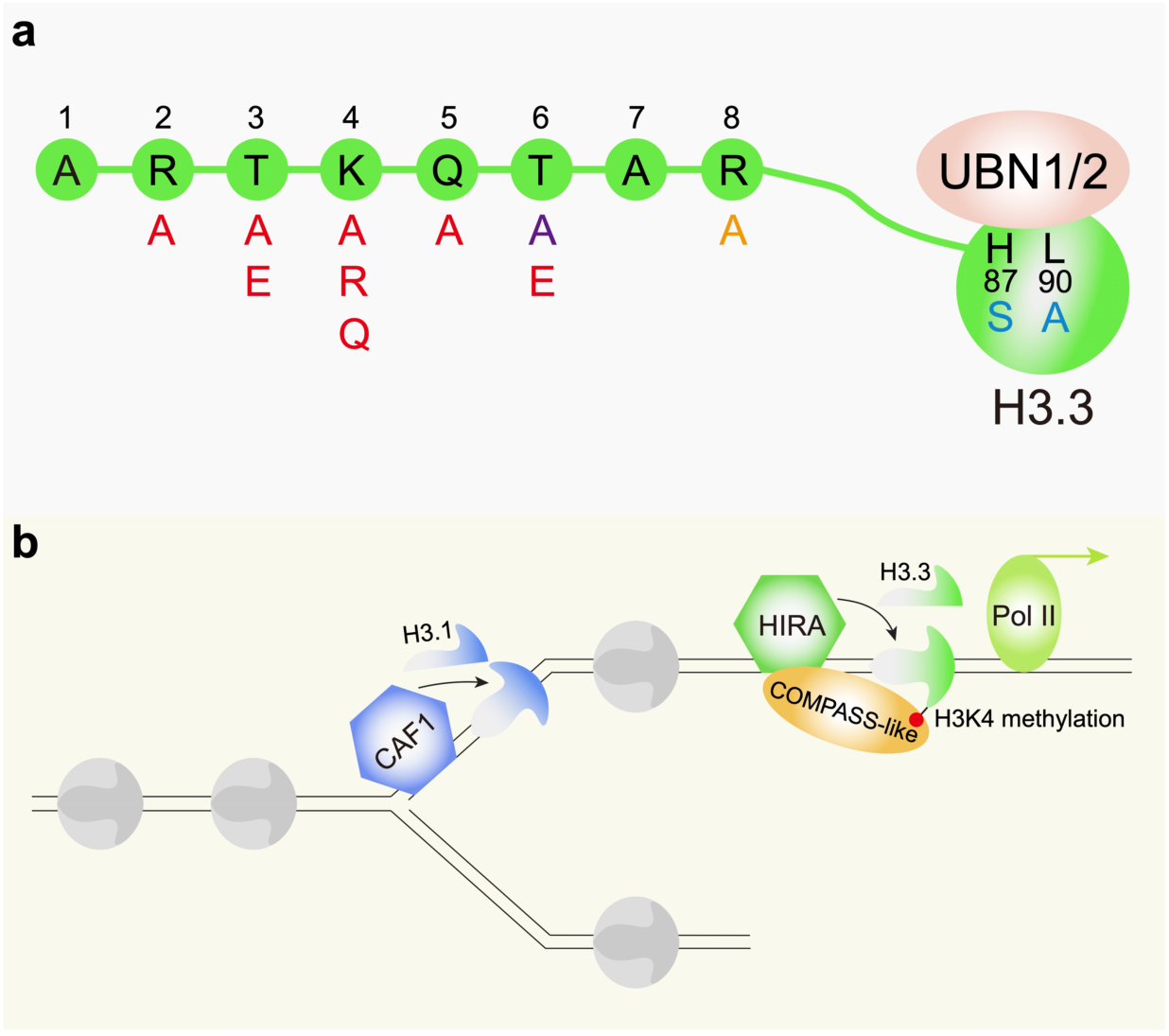
A summary of histone H3.3 mutations studied in this work and a proposed model for the preferential enrichment of H3K4 methylation on H3.3. **a.** R2A, T3A, T3E, K4A, K4R, K4Q, Q5A, and T6E mutations induce similar phenotypes, likely affecting the enrichment of H3K4 methylation and/or its recognition in cis. The T6A mutation results in gain-of-function effects that impair H3K4 methylation both in cis and in trans. R8A mutation induces less severe phenotypes compared to the other mutations and may affect H3.3 function through H3K4 methylation-dependent or independent pathways. The H87S and L90A mutations compromise the recognition of H3.3 by UBN1/2 and the accumulation of H3K4 methylation on H3.3. **b.** The CAF1 complex incorporates H3.1 during DNA replication, while the HIRA complex selectively incorporates H3.3 in a replication-independent manner. The COMPASS-like complex associates with the HIRA complex, which may contribute to the preferential enrichment of H3K4 methylation on H3.3. The closely coordinated deposition of H3.3 and trimethylation of H3.3K4 facilitates gene activation and RNA Pol II elongation.

Although being widely considered as critical for regulating chromatin activity, the causal importance of histone modifications has mainly been inferred from studies disrupting their corresponding enzymes, which may have non-histone substrates or functions unrelated to their enzymatic activities ^5-10^. These issues highlight the need for more direct approaches to address the significance of histone modifications. We found that mutating K4 in H3.3 to A, R or Q all induced similar developmental defects (Figure 1 and Supplemental Figure 2). R is positively charged like K, and Q could mimic acetylation. While a previous study reported that the H3.3K4 regulates H3.3 deposition in mouse embryonic stem cells ^109^, it appears not to be essential for H3.3 deposition in *Arabidopsis* (Supplemental Figure 3). These observations, together with the findings that the *h3.3ko;HTR5 K4A* shows a strong reduction in H3K4 methylation and exhibits comparable phenotypes and transcriptomic changes to the *sdg2* mutant (Figure 2 and 3) ^23,26^, strongly suggest that H3.3K4 is a crucial substrate for SDG2 function.

A complete loss of H3.3 severely impairs plant germination and post-embryonic development ^59^. Introducing K4 mutated H3.3 could partially restore these defects (Figure 1 and Supplemental Figure 2), indicating that K4 only contributes to part of the H3.3 function. Consistently, while H3.3 is required for chromatin accessibility and gene body DNA methylation ^56,59^, loss of H3.3K4 only slightly affects chromatin accessibility and does not impact gene body DNA methylation (Supplemental Figure 4c and 4d). By utilizing *h3.3ko;HTR5 K4A* and the *sdg2* mutant, we reveal that H3K4me3 is involved in de novo gene activation and RNA Pol II elongation but not initiation (Figure 4). This is similar to recent findings in mouse embryonic stem cells, where H3K4me3 facilitates transcriptional elongation by releasing paused Pol II ^83,84^, suggesting a conserved function of H3K4me3 in both animals and plants. Intriguingly, chaperones that mediate H3.3 deposition physically associate with RNA Pol II and transcription elongation factors ^57,110^. This suggests that the incorporation of H3.3 and the deposition of H3K4me3 may be closely coordinated in regulating RNA Pol II elongation (Figure 8b). In addition to H3K4me3, which predominantly relies on SDG2, levels of H3K4me1 and H3K4me2 are significantly reduced in *h3.3ko;HTR5 K4A* as well (Figure 2a), suggesting that H3.3K4 is also critical for other H3K4 methyltransferases responsible for these two modifications ^25,30^. However, the importance of H3K4me1 and H3K4me2 still need to be determined.

Besides the H3.3K4 mutations, R2A, T3A, T3E, Q5A, and T6E mutations in H3.3 also resulted in similar phenotypes (Figure 7b and Supplemental Figure 9 and 12), indicating a close connection between these residues and H3K4 methylation. These findings are in agreement with previous studies in yeast or animal cells, which demonstrate the importance of H3R2, T3, and Q5, and their respective modifications, such as methylation, phosphorylation and serotonylation, in regulating H3K4 methylation or recognition ^90,91,93^. Therefore, these residues may have conserved functions in both animals and plants. Moreover, the R8 residue also contributes to the function of H3.3 but not H3.1, although the phenotype induced by H3.3 R8A mutation is less severe (Figure 7b and Supplemental Figure 9). It remains to investigate whether R8 and its methylation influence H3K4 methylation or if they play independent roles that are important for the H3.3 function.

Mutations of lysine residues in H3 such as K4M, K9M, K27M, and K36M, induce gain-of-function effects that strongly impair the deposition of H3K4, H3K9, H3K27, and H3K36 methylation, respectively ^62-67^. Our results demonstrate that mutating H3.3T6 to A causes similar effects (Figure 7c-7e and Supplemental Figure 13a-13d), suggesting that not only lysine but also other residue mutations could dominant-negatively affect the activity of histone lysine methyltransferases. Interestingly, the T6A mutation-induced dominant-negative effect is specific to H3.3 and not H3.1 (Figure 7c, 7l and 7m), supporting the preferential association between the H3K4 methylation machinery and H3.3. While many missense mutations in histones have been associated with cancer, their significance and specific roles in tumorigenesis remain largely unexplored^111^. Our study has demonstrated that plants can serve as a powerful platform for understanding the function of histones at the single amino acid level.

## Materials and methods

### Plant materials and growth conditions

*h3.1kd* ^11^*, h3.3ko* ^59^ and the *sdg2* mutant (Salk_021008) ^26^ were reported previously. Plants were grown under long day conditions (16h light/8h dark) at 22°C. For high temperature treatment, 3-week-old seedlings were shifted from 22°C to 28°C or kept at 22°C for 4 hours.

### Seed germination test

Seeds were collected at the same time from mother plants grown side by side. After-ripened seeds (seeds harvested and stored at room temperature for over three months) were sown on 1/2 MS medium and stratified at 4°C for three days. Subsequently, seeds were subjected for germination under long days at 22°C. Radicle protrusion was considered as the completion of seed germination. All germination experiments were performed with three biological replicates.

### Plasmid construction for plant transformation

For the *h3.1kd* and *h3.3ko* rescue experiments, the H3.1 coding gene *HTR13*, the H3.3 coding gene *HTR5*, or their mutated forms were cloned into the binary vector pBilligatorR43 as previously described ^11,59^. The *HTR13* coding sequences were additionally mutated, without changing the protein sequences, at the binding site of an artificial microRNA that targets *HTR13* in *h3.1kd* ^11^. For HTR5-GFP and mutated HTR5-GFP constructs, the *HTR5* or mutated *HTR5* genomic sequences, including its promoter, were inserted into pGWB540 ^112^ to fuse with GFP. For HTR13-HA, HTR5-HA and mutated HTR5-HA constructs, the *HTR13*, *HTR5* or mutated *HTR5* genomic sequences and *HTR13* or *HTR5* promoter were inserted into pGWB513 ^112^ to fuse with the 3XHA tag.

### Immunofluorescence

Immunostaining with isolated mature leaf nuclei was performed as previously described ^113^. GFP or H2A.W signals were detected with anti-GFP (Roche, 11814460001) or anti-H2A.W ^72^ antibody, respectively. Images were captured with a Zeiss confocal laser scanning microscope.

### Western blotting

For western blot detection of histone and histone modifications, nuclei extracts from 3-week-old seedlings were transferred to a 0.2 μm nitrocellulose membrane (GE Healthcare) after separated by SDS-PAGE. Proteins were detected with anti-H3K4me1 (Abcam, ab8895), anti-H3K4me2 (Abclonal, A22143), anti-H3K4me3 (Abcam, ab8580), anti-H3K27me3 (Millipore, 07-449), anti-H3K36me3 (Abcam, ab9050), anti-H3 (Abcam, ab1791) or anti-HA (CST, 3724) antibody. The intensity of the protein band was quantified with ImageJ and subsequently normalized to the HA or H3 loading control.

### ChIP-seq

ChIP-seq was performed using 3-week-old seedlings, which were fixed with 1% formaldehyde. To profile H3K4me3 and HTR5/HTR5 K4A-GFP, nuclei were extracted, and mononucleosomes were generated using micrococcal nuclease (NEB, M0247S) digestion. Immunoprecipitation was conducted with anti-H3K4me3 (Abcam, ab8580) or anti-GFP (Thermo Fisher Scientific, A11122) antibody. To profile RNA Pol II Ser5p and RNA Pol II Ser2p, chromatin was sheared to generate 200-400bp fragments and immunoprecipitated with anti-RNA Pol II Ser5p (Abcam, ab5131) or anti-RNA Pol II Ser2p (Abcam, ab5095) antibody as previously described ^114^. Two independent biological replicates were performed for the H3K4me3, RNA Pol II Ser5p and RNA Pol II Ser2p ChIP, and one replicate was performed for the HTR5/HTR5 K4A-GFP ChIP. Mononucleosomes extracted from human HEK293 cells were added as a “spike-in” reference before immunoprecipitation with anti-H3K4me3. The input DNA and immunoprecipitated DNA were subjected to library preparation with VAHTS universal DNA library prep kit for illumina (Vazyme, ND607-02) according to the manufacturer’s instruction. Prepared libraries were sequenced on a NovaSeq 6000 platform and paired-end 150bp reads were generated. Adapter trimming was performed and low quality reads were filtered with fastp version 0.20.1 ^115^. Reads were mapped to the *Arabidopsis* (TAIR10) or human (hg38) genome with Botiew2 version 2.4.2 ^116^. Duplicate reads were filtered using Picard version 2.24.0 MarkDuplicates (https://github.com/broadinstitute/picard). H3K4me3 peaks in WT were called using MACS2 version 2.1.2 with default parameters ^117^. Only peaks identified from both biological replicates were retained. For data visualization, data from two biological replicates were merged, and bigwig coverage files were generated using deepTools utility bamCoverage with a bin size of 10bp ^118^. Spike-in normalization factors were used for the normalization of H3K4me3 ChIP-seq data. Average ChIP-seq profiles were generated using deepTools utility plotProfile.

For the correlation analysis between H3.1, H3.3, H3K4me1, H3K4me2, H3K4me3, H3K27me1 and H3K27me3, as well as the analysis of H3.1 and H3.3 signals over H3K4me1, H3K4me2 or H3K4me3-enriched peaks, relevant ChIP-seq data were obtained from previous publications ^38,119-122^.

### BS-seq

Genomic DNA was extracted from 3-week-old seedlings with Quick-DNA Plant/Seed Miniprep Kit (ZYMO, D6020). 200ng fragmented genomic DNA (200bp-500bp) was used for library preparation with VAHTS universal Pro DNA library prep kit for Illumina (Vazyme, ND608). NEBNext Multiplex Oligos for Illumina (NEB, E7535) were used for adapter ligation. After purification with VAHTS DNA Clean Beads (Vazyme N411), bisulfite conversion was performed with EZ DNA Methylation-Gold Kit (ZYMO, D5005), followed by DNA purification and PCR amplification. Libraries were sequenced with Illumina NovaSeq 6000 to generate paired-end 150bp reads. BS-seq experiments were performed with two independent biological replicates. Adapter trimming was performed and low-quality reads were filtered with fastp version 0.20.1 ^115^. Reads were mapped to the *Arabidopsis* genome (TAIR10) with BS-Seeker2 version 2.1.8 using default parameters ^123^. Duplicated reads were filtered with Picard version 2.24.0 MarkDuplicates (https://github.com/broadinstitute/picard). For data visualization, data from two biological replicates were merged. CG, CHG, and CHH methylation levels were calculated with CGmaptools (version 0.1.2) ^124^. Methylation data were visualized with ggplot2 in R (version 1.1.1106).

### ATAC-seq

Three-week old seedlings were chopped in lysis buffer (15 mM Tris-HCl pH 7.5, 20 mM NaCl, 80 mM KCl, 0.5 mM Spermine, 0.2% Triton X-100, 5 mM β-ME, protease inhibitor cocktail). After being filtered with a 30μm filter (CellTrics), nuclei were stained with DAPI and subjected to fluorescence-activated cell sorting (FACS). 50,000 nuclei per sample were collected and mixed with Tn5 transposase (Mei5bio, MF650-01), after 20 minutes of incubation at 37°C, tagmented DNA was recovered with ChIP DNA Clean & Concentrator kit (ZYMO, D5205). After PCR amplification, sequencing libraries were purified with VAHTS DNA Clean Beads (Vazyme N411) and sequenced with Illumina NovaSeq 6000 to generate paired-end 150bp reads. ATAC-seq experiments were performed with two independent biological replicates. Adapter trimming was performed and low-quality reads were filtered with fastp version 0.20.1 ^115^. Reads were mapped to the *Arabidopsis* genome (TAIR10) with Bowtie2 version 2.4.2 ^116^. Reads mapped to chloroplast and mitochondria genome were removed and duplicated reads were filtered with Picard version 2.24.0 MarkDuplicates (https://github.com/broadinstitute/picard). For data visualization, data from two biological replicates were merged and bigwig coverage files were generated using deepTools utility bamCoverage ^118^ with a bin size of 10bp and normalized with RPKM. Average ATAC-seq profiles were generated using deepTools utility plotProfile.

### RNA-seq

For RNA-seq analysis, total RNA was extracted from 3-week-old seedlings with Minibest plant RNA extraction kit (Takara, 9769) and three independent biological replicates were performed. Sequencing libraries were prepared with the NEBNext Ultra RNA library prep kit for Illumina (NEB, 7530L) according to the manufacturer’s instruction. Prepared libraries were sequenced on a NovaSeq 6000 platform and paired-end 150bp reads were generated. Adapter trimming was performed and low quality reads were filtered with fastp version 0.20.1 ^115^. Reads were aligned to the *Arabidopsis* genome (TAIR10) using Hisat2 version 2.1.10 ^125^. Reads per gene were counted by HTseq version 0.11.2 ^126^. Transcripts per million (TPM) values were generated using R. Differential gene expression analysis was performed using DESeq2 version 1.26.0 ^127^. Genes were considered as differentially expressed in RNA-seq if they exhibited a more than two-fold change in expression and had a *P* adjust value<0.05. Gene ontology analysis was performed with DAVID (https://david.ncifcrf.gov/) ^128^. IR analysis was performed using IRFinder (version 1.3.1) ^129^. The IR ratio was calculated as intronic abundance divided by the sum of intronic abundance and normal splicing abundance ^129^. The significance of differential IR between two samples with three biological replicates each was tested with the generalized linear model in DESeq2 (version 1.26.0). Introns with IR ratio change more than 0.1 and P adjust value less than 0.05 were considered as differentially spliced.

### RT-qPCR

Total RNA was extracted from 3-week-old seedlings with Minibest plant RNA extraction kit (Takara, 9769) and three independent biological replicates were performed. Reverse transcription was performed using HiScript III 1^st^ Strand cDNA Synthesis Kit (Vazyme, R312-02). Real-time quantitative PCR was conducted on an Applied Biosystems QuantStudio 6 Flex Real-Time PCR System using ChamQ Universal SYBR qPCR Master Mix (Vazyme, Q711-02). *ACT7* was used as an endogenous control for normalization. Primers used for amplification are listed in Supplemental table 1.

### ChIP-qPCR

Three-week-old seedlings were shifted from 22°C to 28°C or kept at 22°C for 4 hours. After fixed with 1% formaldehyde, chromatin was sheared to generate 200-400bp fragments and immunoprecipitated with anti-H3K4me3 (Abcam, ab8580), anti-RNA Pol II Ser5p (Abcam, ab5131) or anti-RNA Pol II Ser2p (Abcam, ab5095) antibody as previously described ^114^. The amount of immunoprecipitated DNA was quantified by real-time PCR. Three independent biological replicates were performed. Primers used for amplification are listed in Supplemental table 1.

### Mononucleosome reconstitution

The full-length coding sequences of *Arabidopsis* H2A (HTA10, *AT1G51060*), H2B (HTB1, *AT1G07790*), H3.1 (HTR2, *AT1G09200*), H3.3 (HTR8, *AT5G10980*) and H4 (*AT2G28740*) were cloned into pET28a vector and histone proteins were expressed using *E.coli* BL21 (DE3) by IPTG induction. Cell pellets were collected and sonicated. After centrifugation, the inclusion bodies were resuspended in unfolding buffer (7 M guanidinium HCl, 20 mM Tris-HCl pH 7.5, 10 mM DTT) and rotated for 1 hour at room temperature, and the supernatant containing histone was collected after centrifugation.

In vitro nucleosome assembly were performed as described previously ^130^. In brief, equimolar H2A, H2B, H3.1 or H3.3 and H4 histones were mixed and dialyzed against 2 L refolding buffer (2 M NaCl, 10 mM Tris-HCl pH 7.5, 1 mM EDTA, 5 mM 2-Mercaptoethanol) overnight at 4 °C with two changes of refolding buffer. The histone octamers were purified with Superdex 200 increase 10/300 GL column (GE). Mononucleosomes were assembled using the salt-dialysis method. Octamers and single 601 DNA (225 bp amplified by PCR) were mixed in the assembly buffer (10 mM Tris-HCl pH 8.0, 1 mM EDTA, 2 M NaCl) and then the mixture was dialyzed against 450 ml assembly buffer. Subsequently, the concentration of NaCl was gradually reduced to 0.6 M by pumping in TE buffer (10 mM Tris-HCl pH 8.0, 1 mM EDTA). Finally, mononucleosomes were dialyzed with TE buffer overnight.

### *In vitro* histone methyltransferase assay

To produce GST-SDG2set and MBP-ATXR6set, the SET domain coding sequences of SDG2 or ATXR6 were cloned into pGEX-5X-2 or pMAL-C5X, respectively. Proteins were expressed using *E.coli* BL21 (DE3) by IPTG induction. Cell pellets were resuspended with lysis buffer (150 mM NaCl, 20 mM Tris-HCl pH 8.0, 10% Glycerol) and sonicated. After centrifugation, the supernatant was mixed with glutathione-agarose resin (GE Healthcare 17075601) or amylose resin (NEB, E8021) and rotated at 4°C overnight. Beads were washed three times with lysis buffer and once with elution buffer (10 mM Tris-HCl pH 8.0, 1mM EDTA), and GST-SDG2set or MBP-ATXR6set protein were then eluted with elution buffer containing 20 mM reduced glutathione or 10 mM maltose.

For *in vitro* histone methyltransferase assay, 2 μg mononucleosomes were mixed with 2 μg GST-SDG2set or 5ug MBP-ATXR6set in 30 μl HMT buffer (50 mM Tris-HCl pH 8.5, 1 mM MgCl_2_, 4 mM DTT) supplemented with 80 μM S-adenosylmethionine (NEB, B9003S) at 30°C overnight. Reactions were stopped by adding the SDS loading buffer and followed by incubating at 100°C for 5 minutes. After electrophoresis, proteins were transferred to a 0.2 μm PVDF membrane (Millipore), histone modifications were detected with anti-H3K4me1 (Abcam, ab8895), anti-H3K4me2 (Abclonal, A22143), anti-H3K4me3 (Abcam, ab8580) or anti-H3K27me1 (Millipore, 07-448) antibody. Protein loading was detected by coomassie blue staining of the membrane.

### Mononucleosome pull-down assay

Mononucleosomes pull-down was performed as previously described with minor modifications ^131^. Mononucleosome was assembled with histone octamers and biotin-labeled 601 DNA that was prepared by PCR amplification with biotin-labeled oligos. 2 μg biotin-labeled mononucleosomes and 20 μl Streptavidin C1 beads (ThermoFisher, 65001) were mixed in BC300 buffer (20 mM Tris–HCl pH 7.5, 10% glycerol, 300 mM NaCl, 0.1% NP-40, 500 μg/ml BSA), and then incubated with GST-SDG2set overnight at 4°C. Beads were washed with wash buffer (20 mM Tris–HCl pH 7.5, 10% glycerol, 300 mM NaCl, 0.1% NP-40) for three times. Proteins retained on the beads were eluted by boiling with SDS loading buffer, separated by SDS-PAGE, and detected by anti-GST (Easybio, BE2013), anti-H3 (Abcam, ab1791) or anti-H2B (Abcam, ab1790) antibody.

### Pull-down assay

To produce MBP-HIRA, full-length coding sequence of *HIRA* was cloned into pMAL-C5X. To produce GST-UBN1 or GST-UBN2, full-length coding sequences of *UBN1* or *UBN2* was cloned into pGEX-5X-2. To produce His-H3.1, His-H3.3 or His-RBL, full-length coding sequences of *HTR13*, *HTR5* or *RBL* was cloned into pRSETA. Proteins were expressed using *E.coli* BL21 (DE3) by IPTG induction. After incubating with His-tagged proteins in pull-down buffer (50 mM Tris pH 7.5, 150 mM NaCl, 1 mM EDTA, 0.5% Nonidet P-40, protease inhibitor cocktail) at 4°C overnight, GST-tagged or MBP-tagged proteins were captured by glutathione-agarose resin (GE Healthcare, 17075601) or amylose resin (NEB, E8021) respectively. Beads were washed four times with pull-down buffer and proteins retained on the beads were eluted by boiling with SDS loading buffer, separated by SDS-PAGE, and detected with anti-GST (Easybio, BE2013), anti-MBP (NEB, E8032S) or anti-His (CWBio, CW0286) antibody.

### BiFC

The BiFC experiment was performed as previously described ^132^. The full-length coding sequences of *HTR5*, *HTR5 H87S&L90A*, *HTR13*, *UBN1*, *UBN2*, *HIRA* or *RBL* without stop codon were cloned into pEarleyGate201-YN or pEarleyGate202-YC vector to fuse with nYFP (YN) or cYFP (YC) respectively. In each transformation, same amount of agrobacterium containing YN, YC and mRFP-AHL22 were used for co-infiltration into *N. benthamiana* plant leaves. YFP and RFP signals were observed 2 days after infiltration using a Zeiss confocal laser-scanning microscope with the same setting for each transformation. The intensity of YFP and RFP signals in nuclei were quantified with ImageJ.

### Co-IP assay

To express tag-fused RBL and HIRA, their genomic sequences including their respective promoters were cloned into pGWB510 and pGWB516 respectively^112^. To test the RBL-HIRA, RBL-H3.1 and RBL-H3.3 protein interactions, *Arabidopsis* plant materials were generated by crossing *RBL-FLAG* with *HIRA-Myc*, *HTR13-HA* or *HTR5-HA* transgenic lines. Proteins were extracted with Co-IP buffer (50 mM HEPES pH 7.5, 150 mM NaCl, 1 mM EDTA, 1mM DTT, 0.3% Triton X-100, protease inhibitor cocktail). After centrifugation, the supernatant was incubated with anti-Myc (Thermo scientific, 88842) or anti-FLAG (abmart, M20018S) beads. Tagged proteins were detected with anti-Myc (Easybio, BE2011), anti-FLAG (Proteintech, 20543-1-AP) or anti-HA (Proteintech, 51064-2-AP) antibody.

### Dot blot assay

WT or mutated lysine 4 trimethylated H3 (1-20aa) peptides were synthesized by GenScript Biotech Corporation. Peptides were dotted in titration onto a nitrocellulose membrane (GE Healthcare) and allowed to air dried for approximately 30 min to 1 h. The peptides were detected with anti-H3K4me3 (Abcam, ab8580) antibody, and peptides loading was assessed by staining the membrane with Ponceau S.

### Protein structure predication and docking

The protein structure of SDG2 SET domain was predicted with AlphaFold 2 using ColabFold v1.5.2(https://github.com/sokrypton/ColabFold) ^133^. PDB files for histone H3 peptides (1-15aa) were designed by PYMOL (http://www.pymol.org/). Protein-protein docking was performed with ClusPro (https://cluspro.bu.edu/) ^134^. The structural figures were created using ChimeraX (https://www.cgl.ucsf.edu/chimerax/).

### Statistical analysis

The significance of differences was determined with two-tailed Student’s *t*-test (*, *P* < 0.05; **, *P* < 0.01; ns, not significant) or one-way ANOVA with Tukey’s test. The significance of overlap between two sets of genes was calculated with hypergeometric test.

## Data availability

The datasets generated in this study are available in the GEO. The private access tokens are GSE274521: sjwjccomvrkpvqp, GSE274522: utihikkevlubvyx, GSE274523: elursuewdxkrlmt, GSE274524: onwngoeshvqjdmb, GSE274525: cfktkascvngrpql, and GSE274526: shqvemsgrxkffmj.

## Author contributions

M.X., L.M. and D.J. designed experiments, M.X., L.M., X.L. and D.J. performed experiments, M.X., L.M., H.Z. and Q.L. analysed data, D.J. wrote the manuscript with the help from M.X. and L.M.

## Acknowledgements

We thank Dr. Cuifang Liu for valuable suggestions regarding in vitro assembly of nucleosomes, and Dr. Fengyue Zhao for providing plasmids. This work was supported by the National Key R&D Program of China (2019YFA0903903), the Strategic Priority Research Program of the Chinese Academy of Sciences (Precision Seed Design and Breeding, XDA24020303), the National Natural Science Foundation of China (32170545) and the intramural research support from Temasek Life Sciences Laboratory.

## Notes

### Competing Interest Statement

The authors have declared no competing interest.

