## Supplemental Figures and Tables for "Single amino acid mutations in histone H3.3 illuminate the functional significance of H3K4 methylation in plants"

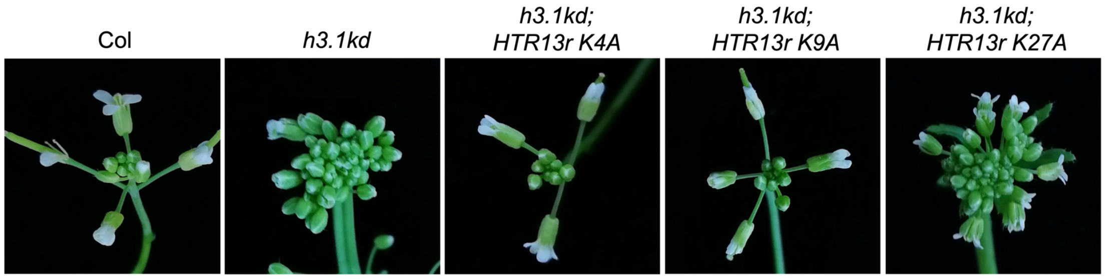

**Supplemental Figure 1. Inflorescence developmental phenotypes of the indicated *h3.1kd* rescue lines**

*HTR13r* represents the mutated *HTR13* at the binding site of an artificial microRNA targeting *HTR13* in *h3.1kd*, without altering the protein sequences.

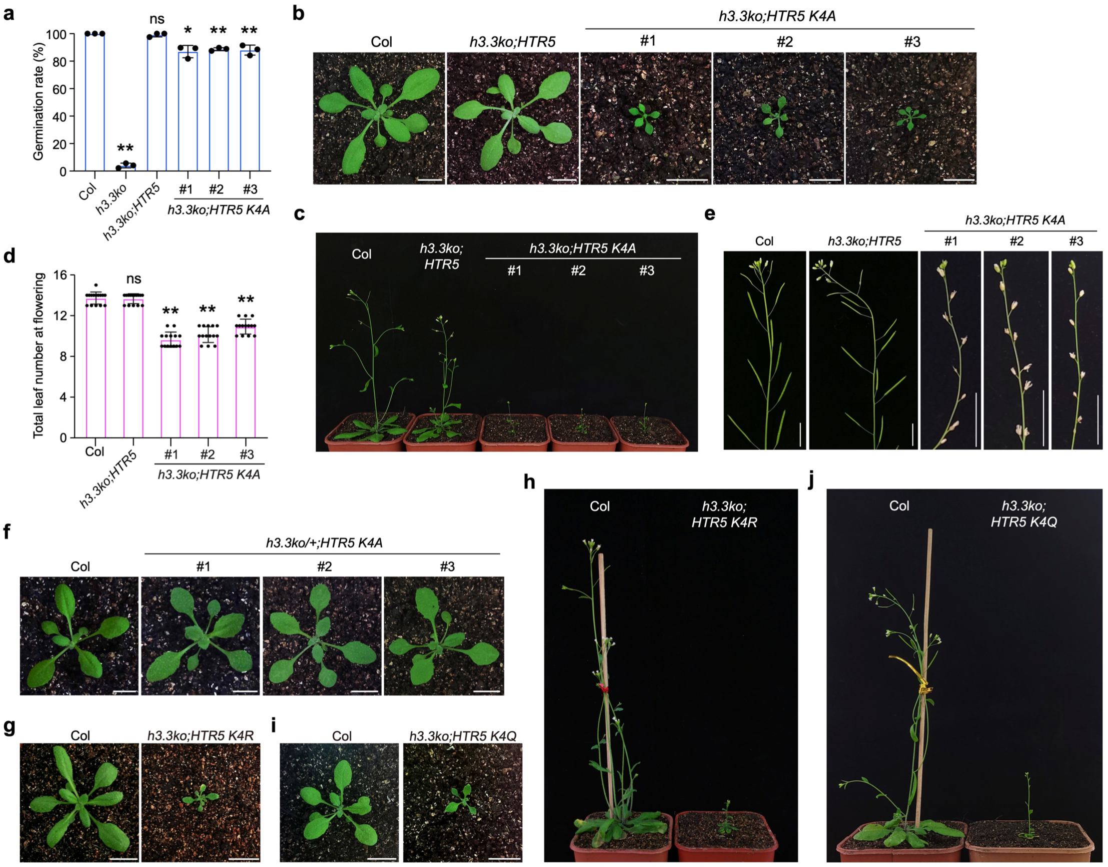

**Supplemental Figure 2. Developmental phenotypes of *h3.3ko;HTR5 K4A*, *h3.3ko;HTR5 K4R* and *h3.3ko;HTR5 K4Q***

**a.** Seed germination rates of the indicated lines after imbibed for 7 days. Values are means ± SD of three biological replicates. At least 32 seeds were analyzed per replicate. Statistical significance relative to Col was determined by two-tailed Student's *t*-test (*, *P* < 0.05; **, *P* < 0.01; ns, not significant).

**b and c.** Developmental phenotypes of the indicated lines at the vegetative stage (b) and reproductive stage (c). Three independent representative transgenic lines are shown. Scale bars, 1 cm.

**d.** Total number of primary rosette and cauline leaves at flowering for the indicated lines. 14-15 plants were scored for each line. Values are means ± SD. Statistical significance relative to Col was determined by two-tailed Student's *t*-test (**, *P* < 0.01; ns, not significant).

**e.** Silique development phenotypes of the indicated lines. Three independent representative transgenic lines are shown. Scale bars, 1 cm.

**f.** Developmental phenotypes of transgenic lines expressing *HTR5 K4A* in the *h3.3ko/+* (*htr4/htr4;htr5/htr5;htr8/+*) background. Three independent representative transgenic lines are shown. Scale bars, 1 cm.

**g and h.** Developmental phenotypes of Col and *h3.3ko;HTR5 K4R* at the vegetative stage (g) and reproductive stage (h). A representative line is shown. Scale bars, 1 cm.

**i and j.** Developmental phenotypes of Col and *h3.3ko;HTR5 K4Q* at the vegetative stage (i) and reproductive stage (Q). A representative line is shown. Scale bars, 1 cm.

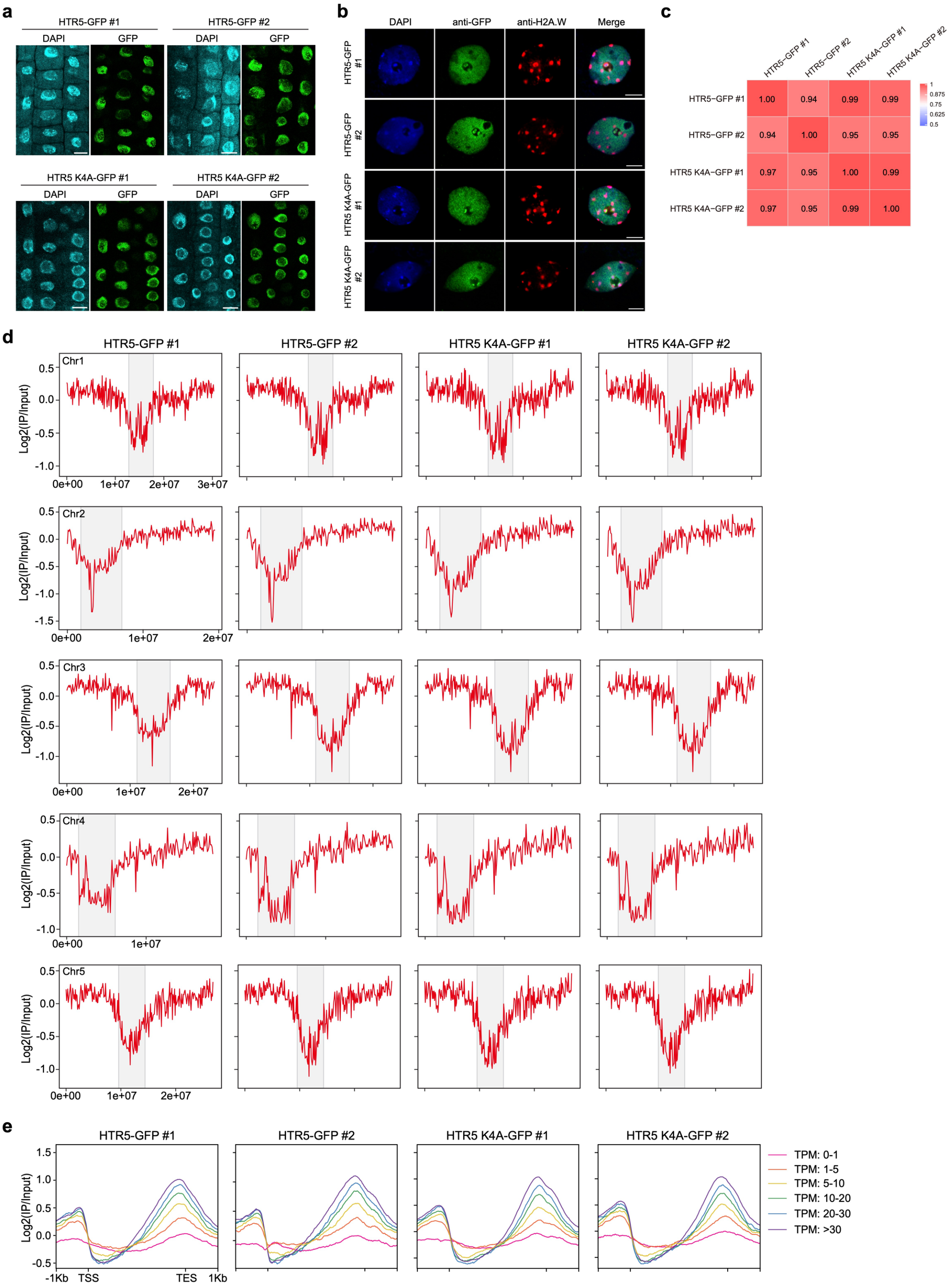

**Supplemental Figure 3. K4 is not required for H3.3 distribution**

**a.** Nuclei localization of GFP-fused H3.3 (HTR5) and H3.3 K4A (HTR5 K4A) in root cells. Nuclei were labeled by 4',6-diamidino-2-phenylindole (DAPI) staining. Results from two independent transgenic lines are shown. Scale bars, 10 μm.

**b.** Chromatin localization of GFP-fused HTR5 and HTR5 K4A as detected by immunofluorescence in leaf nuclei. Condensed chromocenters were immunostained with the heterochromatin mark H2A.W. Results from two independent transgenic lines are shown. Scale bars, 5 μm.

**c.** Heat map showing the Pearson correlation between genome-wide distributions of GFP-fused HTR5 and HTR5 K4A in two independent transgenic lines profiled by ChIP-seq.

**d.** HTR5-GFP and HTR5 K4A-GFP ChIP-seq signals over *Arabidopsis* chromosomes. Signals were calculated in 100kb bins. Pericentromeric heterochromatin regions are indicated with grey shading.

**e.** Metaplot of HTR5-GFP and HTR5 K4A-GFP ChIP-seq signals over genes grouped according to their transcript levels.

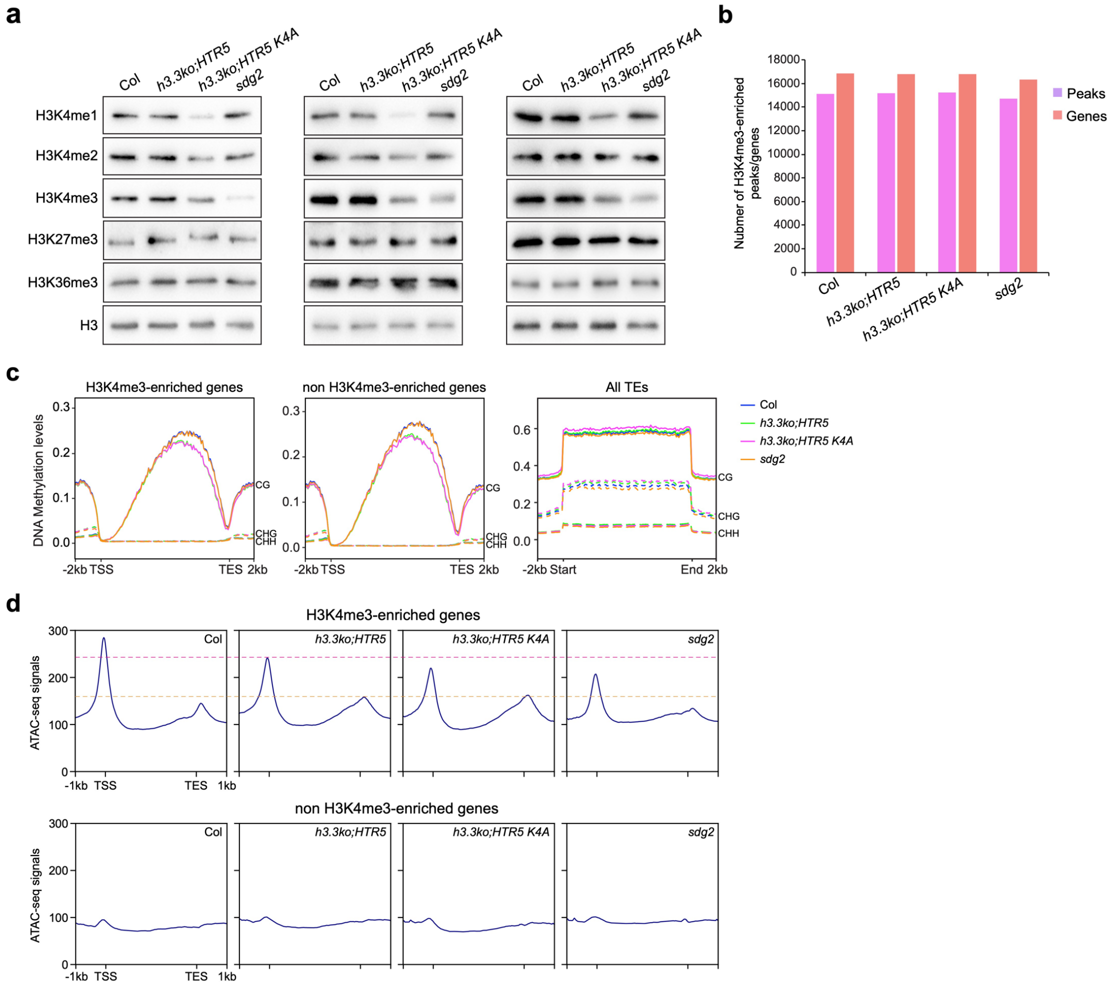

**Supplemental Figure 4. Analysis of histone modifications, DNA methylation and chromatin accessibility in Col, *h3.3ko;HTR5*, *h3.3ko;HTR5 K4A* and *sdg2***

**a.** Histone modification levels in Col, *h3.3ko;HTR5*, *h3.3ko;HTR5 K4A* and *sdg2* determined by western blotting. H3 was employed as a loading control. Results from three biological replicates are presented.

**b.** Number of H3K4me3-enriched peaks and genes identified in Col, *h3.3ko;HTR5*, *h3.3ko;HTR5 K4A* and *sdg2*.

**c.** Metaplot of DNA methylation levels in Col, *h3.3ko;HTR5*, *h3.3ko;HTR5 K4A* and *sdg2* over H3K4me3-enriched genes, non H3K4me3-enriched genes in WT Col, and all TEs.

**d.** Metaplot of ATAC-seq signals in Col, *h3.3ko;HTR5*, *h3.3ko;HTR5 K4A* and *sdg2* over H3K4me3-enriched genes and non H3K4me3-enriched genes in WT Col.

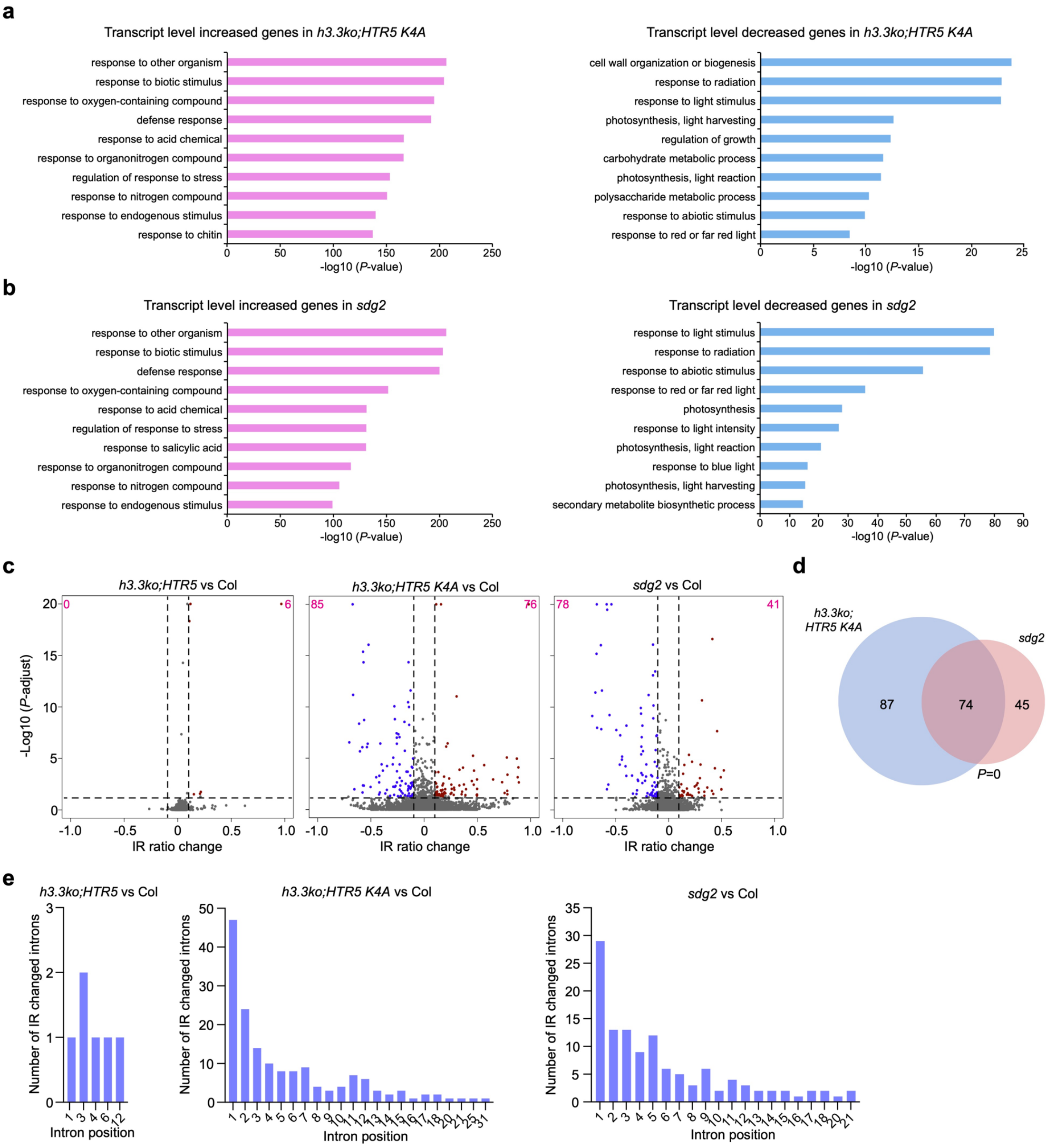

**Supplemental Figure 5. Analysis of transcriptome changes in *h3.3ko;HTR5 K4A* and *sdg2***

**a and b.** Gene ontology (GO) analysis of misexpressed genes in *h3.3ko;HTR5 K4A* (a) or *sdg2* (b). Top 10 representative terms are listed and ranked by *P* value.

**c.** Volcano plot of IR changes determined by RNA-sequencing. The y-axis values correspond to -log$_{10}$ (*P*-adjust), and the x-axis values correspond to IR ratio change. Introns with IR ratio change more than 0.1 and *P*-adjust less than 0.05 were considered differentially spliced. The numbers of IR ratio increased and decreased introns are indicated at the top right and left corners, respectively.

**d.** Venn diagrams of IR ratio changed introns in *h3.3ko;HTR5 K4A* and *sdg2* compared with Col.

**e.** Distributions of IR changed introns in *h3.3ko;HTR5*, *h3.3ko;HTR5 K4A* and *sdg2* compared with Col.

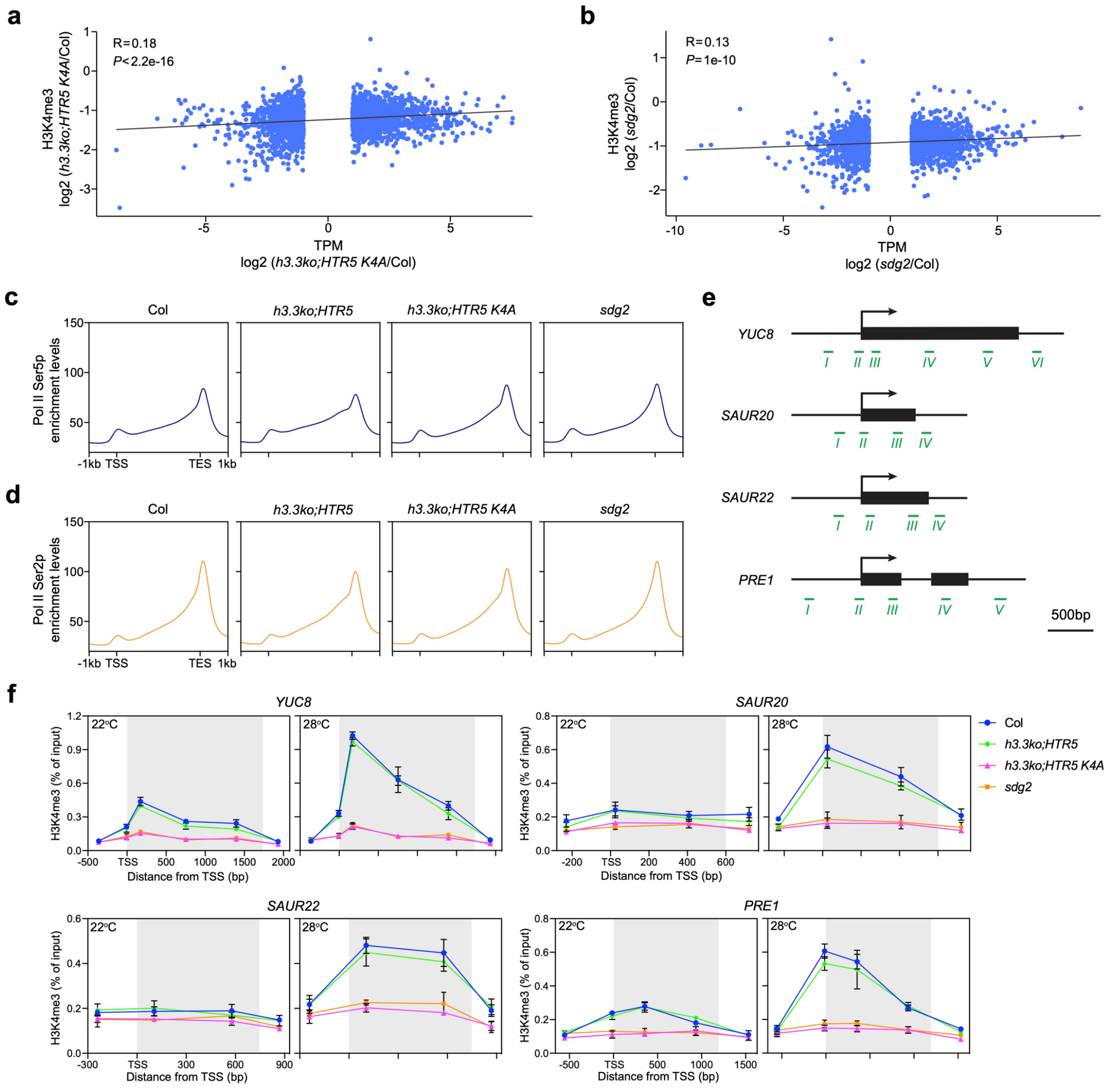

**Supplemental Figure 6. Analysis of gene expression, RNA Pol II enrichment and H3K4me3**

**a and b.** Correlation between changes in transcript levels of significantly misexpressed genes and their H3K4me3 levels (from TSS to 500bp downstream) in *h3.3ko;HTR5 K4A* (a) or *sdg2* (b) compared to Col.

**c and d.** Metaplot of Pol II Ser5p (c) and Pol II Ser2p (d) ChIP-seq signals in Col, *h3.3ko;HTR5*, *h3.3ko;HTR5 K4A* and *sdg2* over H3K4me3-enriched genes in WT Col.

**e.** Schematic structures of *YUC8*, *SAUR20*, *SAUR22*, and *PRE1*. Arrows indicate transcription start sites, and green lines indicate regions examined by ChIP-qPCR.

**f.** H3K4me3 enrichment levels across *YUC8*, *SAUR20*, *SAUR22* and *PRE1* determined by ChIP-qPCR at 22°C and 28°C. Values are means ± SD of three biological replicates. Genic regions are indicated with grey shading.

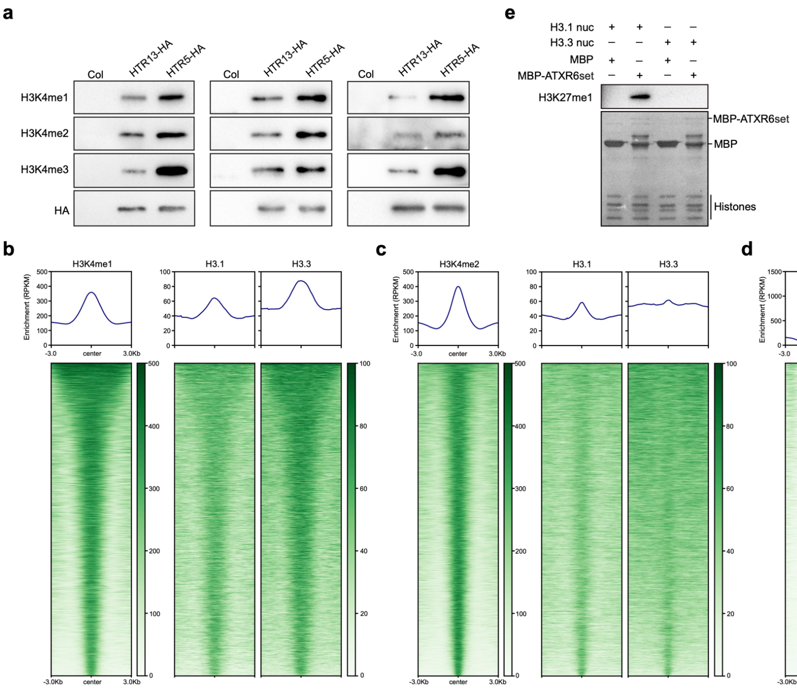

**Supplemental Figure 7. H3.3 positively correlates with H3K4 methylation**

**a.** H3K4 methylation levels on exogenous H3.1 (HTR13) and H3.3 (HTR5) determined by western blotting. Results from three biological replicates are presented.

**b-d.** Metaplot and heatmap of H3.1 and H3.3 ChIP-seq signals over H3K4me1 (b), H3K4me2 (c) and H3K4me3 (d)-enriched peaks.

**e.** *In vitro* methyltransferase assay showing the activity of ATXR6 SET domain (ATXR6set) on nucleosomes containing H3.1 or H3.3. The levels of H3K27me1 in *in vitro* methyltransferase assay products were determined by western blotting.

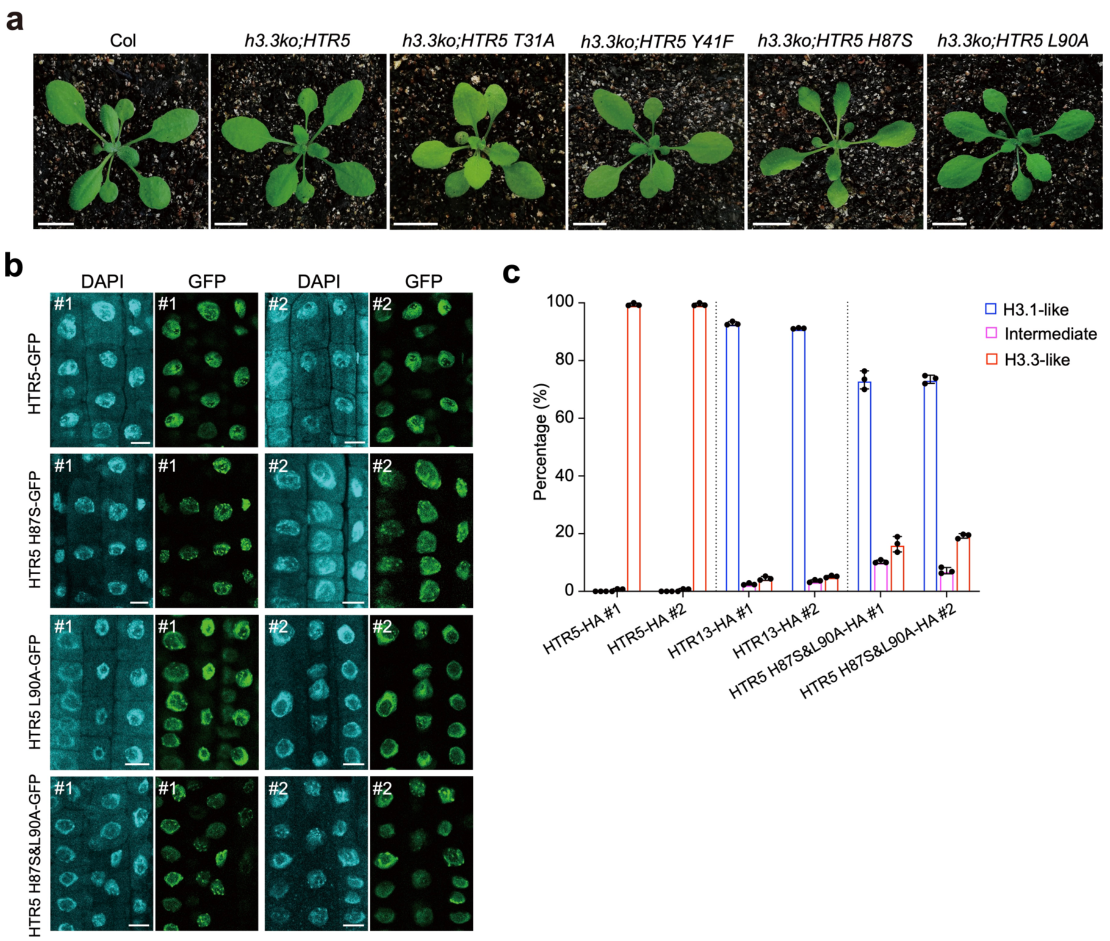

**Supplemental Figure 8. Identification of amino acids required for H3.3 distribution**

**a.** Developmental phenotypes of the indicated lines. A representative line is shown. Scale bars, 1 cm.

**b.** Nuclei localization of GFP-fused H3.3 (HTR5), H3.3 H87S (HTR5 H87S), H3.3 L90A (HTR5 L90A) and H3.3 H87S&L90A (HTR5 H87S&L90A) in root cells. Nuclei were labeled by DAPI staining. Results from two independent transgenic lines are shown. Scale bars, 10 µm.

**c.** Percentages of nuclei showing H3.1-like, intermediate and H3.3-like distribution patterns for the indicated proteins. Results from two independent transgenic lines are shown. Values are means ± SD of three biological replicates. At least 105 nuclei were analyzed per replicate.

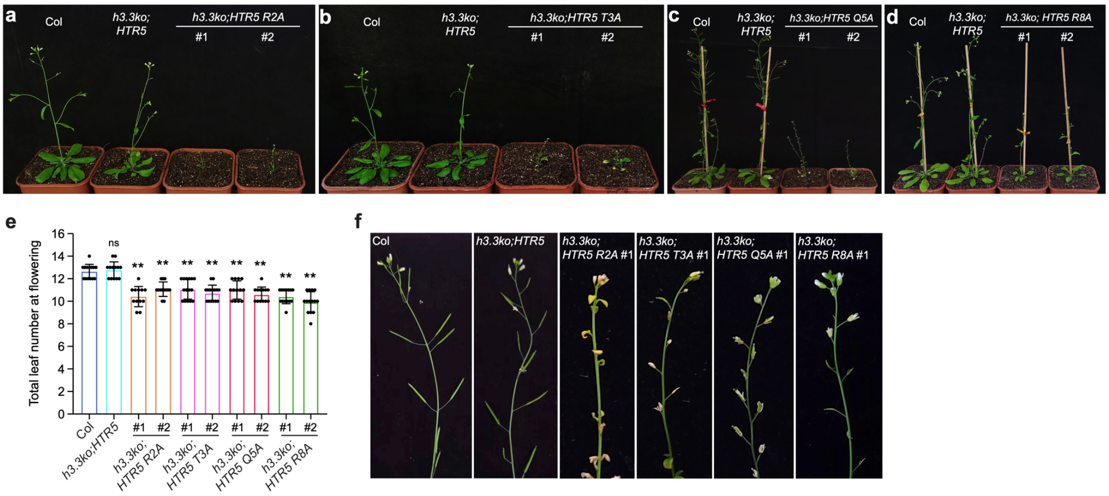

**Supplemental Figure 9. Developmental phenotypes of *h3.3ko;HTR5 R2A*, *h3.3ko;HTR5 T3A*, *h3.3ko;HTR5 Q5A* and *h3.3ko;HTR5 R8A***

**a-d.** Developmental phenotypes of *h3.3ko;HTR5 R2A* (a), *h3.3ko;HTR5 T3A* (b), *h3.3ko;HTR5 Q5A* (c) and *h3.3ko;HTR5 R8A* (d) at the reproductive stage. Two independent representative transgenic lines are shown.

**e.** Total number of primary rosette and cauline leaves at flowering for the indicated lines. 13-15 plants were scored for each line. Values are means ± SD. Statistical significance relative to Col was determined by two-tailed Student's *t*-test (**, $P < 0.01$; ns, not significant).

**f.** Silique developmental phenotypes of the indicated lines.

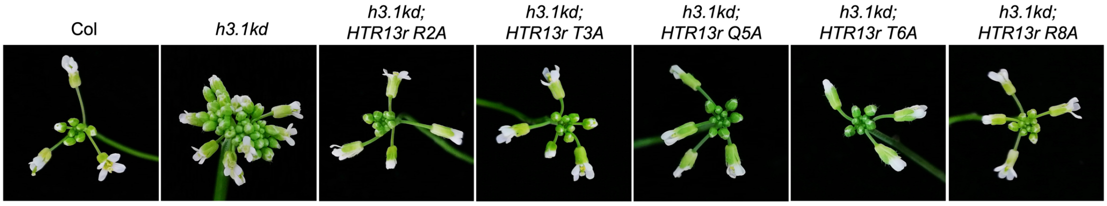

**Supplemental Figure 10. Inflorescence developmental phenotypes of the indicated *h3.1kd* rescue lines**

*HTR13r* represents the mutated *HTR13* at the binding site of an artificial microRNA targeting *HTR13* in *h3.1kd*, without altering the protein sequences.

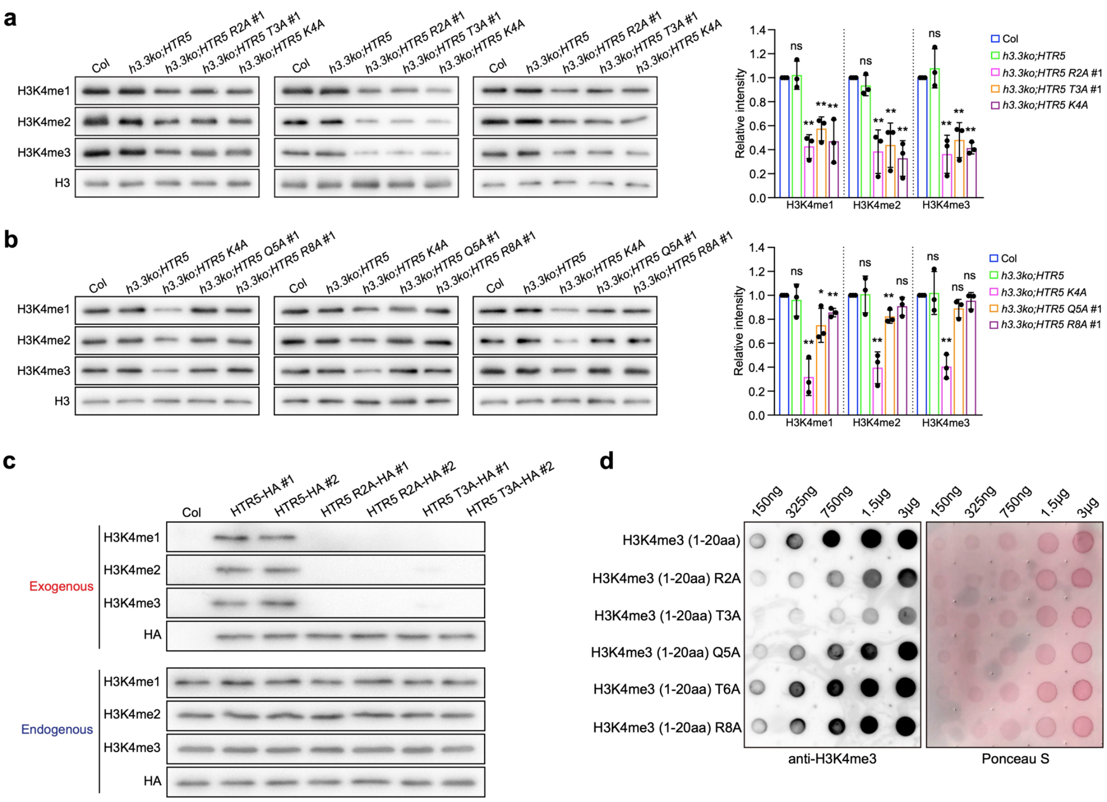

**Supplemental Figure 11. H3K4 methylation changes induced by H3.3 R2A, T3A, Q5A and R8A mutations**

**a and b.** Histone modification levels in the indicated lines determined by western blotting. H3 was employed as a loading control. Results from three biological replicates are presented. The bar charts represent quantification of western blot signals from three biological replicates. Values are means ± SD. Statistical significance relative to Col was determined by two-tailed Student's *t*-test (*, $P < 0.05$; **, $P < 0.01$; ns, not significant).

**c.** H3K4 methylation levels on exogenous H3.3 (HTR5), H3.3 R2A (HTR5 R2A), H3.3 T3A (HTR5 T3A), and endogenous H3 determined by western blotting. Results from two independent transgenic lines are shown.

**d.** Dot blot assay testing the antibody affinity for synthesized H3 peptides (the N-terminal 1-20 amino acids) carrying H3K4me3 and the indicated mutations.

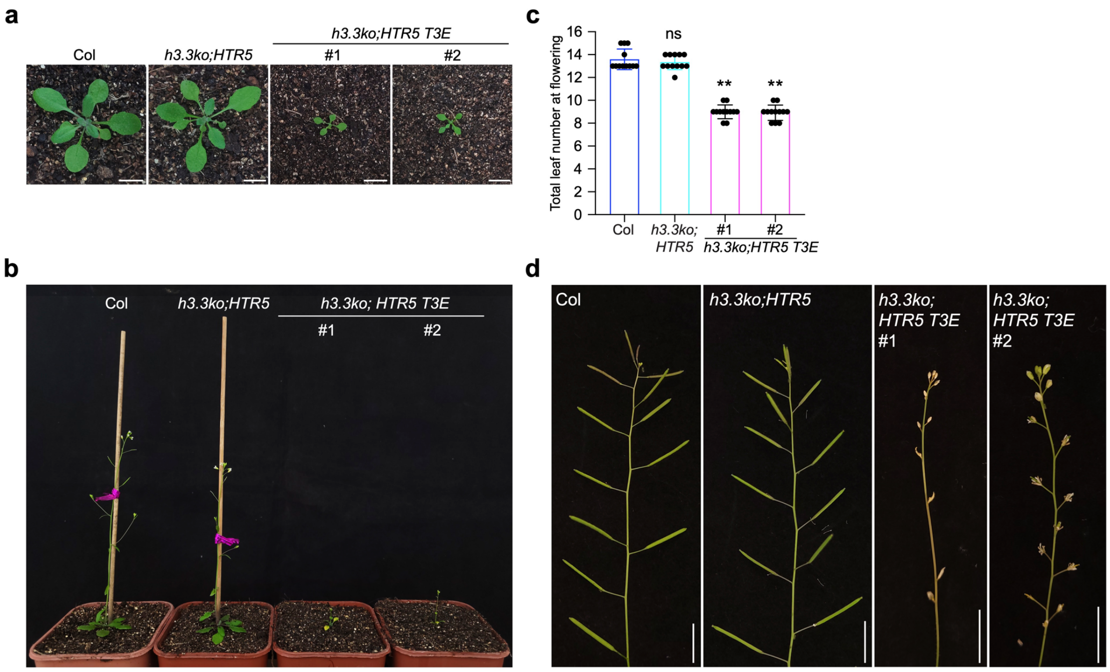

**Supplemental Figure 12. Developmental phenotypes of *h3.3ko;HTR5 T3E***

**a and b.** Developmental phenotypes of the indicated lines at the vegetative stage (a) and reproductive stage (b). Two independent representative transgenic lines are shown. Scale bars, 1 cm.

**c.** Total number of primary rosette and cauline leaves at flowering for the indicated lines. 12 plants were scored for each line. Values are means ± SD. Statistical significance relative to Col was determined by two-tailed Student's *t*-test (**, $P < 0.01$; ns, not significant).

**d.** Silique developmental phenotypes of the indicated lines. Two independent representative transgenic lines are shown. Scale bars, 1 cm.

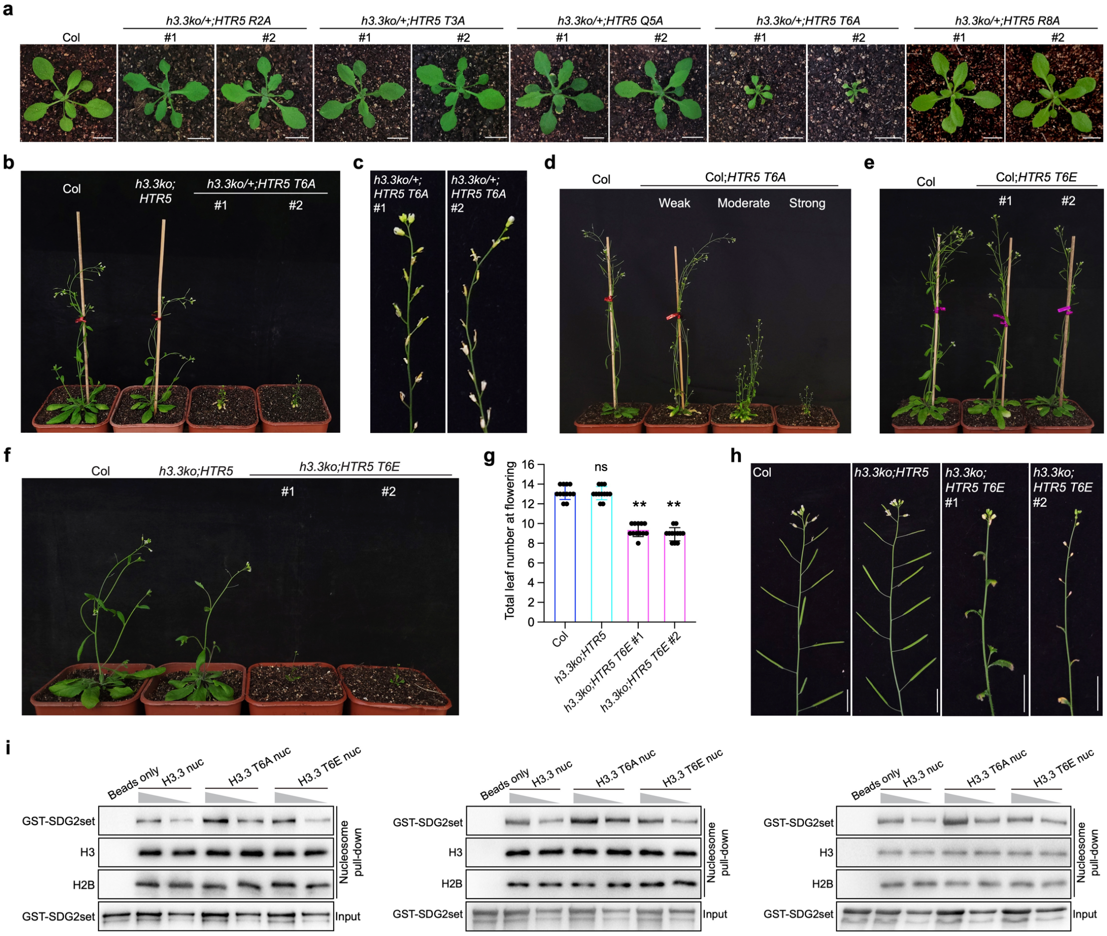

**Supplemental Figure 13. H3.3 T6A or H3.3 T6E mutation-induced phenotypes**

**a.** Developmental phenotypes of transgenic lines expressing *HTR5 R2A*, *HTR5 T3A*, *HTR5 Q5A*, *HTR5 T6A*, or *HTR5 R8A* in the *h3.3ko/+* (*htr4/htr4;htr5/htr5;htr8/+*) background at the vegetative stage. Two independent representative transgenic lines are shown. Scale bars, 1 cm.

**b.** Developmental phenotypes of transgenic lines expressing *HTR5 T6A* in the *h3.3ko/+* (*htr4/htr4;htr5/htr5;htr8/+*) background at the reproductive stage. Two independent representative transgenic lines are shown.

**c.** Silique development phenotypes of transgenic lines expressing *HTR5 T6A* in the *h3.3ko/+* (*htr4/htr4;htr5/htr5;htr8/+*) background. Two independent representative transgenic lines are shown.

**d.** Developmental phenotypes of T1 transgenic lines expressing *HTR5 T6A* in the Col background at the reproductive stage. Representative plants showing weak, moderate and strong phenotypes are shown.

**e and f.** Developmental phenotypes of transgenic lines expressing *HTR5 T6E* in the Col (e) or *h3.3ko* (f) at the reproductive stage. Two independent representative transgenic lines are shown.

**g.** Total number of primary rosette and cauline leaves at flowering for the indicated lines. 12 plants were scored for each line. Values are means ± SD. Statistical significance relative to Col was determined by two-tailed Student's *t*-test (**, $P < 0.01$; ns, not significant).

**h.** Silique development phenotypes of the indicated lines. Two independent representative transgenic lines are shown. Scale bars, 1 cm.

**i.** Mononucleosome pull-down assay of nucleosome containing H3.3, H3.3 T6A or H3.3 T6E incubated with SDG2 SET domain (SDG2set) proteins. Results from three independent replicates are presented.

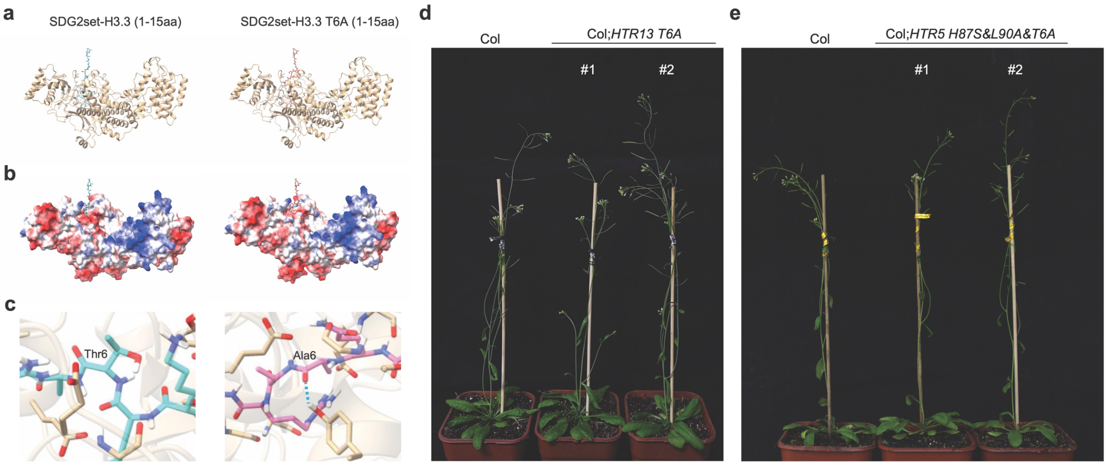

**Supplemental Figure 14. Structural predication and phenotypes of Col plants expressing *HTR13 T6A* or *HTR5 H87S&L90A&T6A***

**a-c.** Predicated structured of SDG2 SET domain (SDG2set) bound to H3.3 or H3.3 T6A peptide. The hydrogen bonding is indicated by a blue dashed line (c). **d and e.** Developmental phenotypes of transgenic lines expressing *HTR13 T6A* (d) or *HTR5 H87S&L90A&T6A* (e) in the Col background at the reproductive stage. Two independent representative transgenic lines are shown.

**Supplemental Table 1. Primers used for RT-qPCR and ChIP-qPCR**

| Experiment | Primer (5' to 3') |
|---|---|
| **RT-qPCR** | |
| ***HTR5*** | TCGTAAGTCTACTGGAGGAAAG |
| | ACGCAAGTCAGTCTTGAAATCC |
| ***YUC8*** | GACTGCTCGGTTCGATGAGAC |
| | TGAATCACCTCACCGGAAAAC |
| ***SAUR20*** | CCTTGTCATGAAGATACGTTCATC |
| | CACAAAAATGGCATCCATTCTCTAAAC |
| ***SAUR22*** | GACAAATAGAGAATTATAAATGGCTCTG |
| | ATGAATTAAGTCTATATCTAACTCGGAAA |
| ***PRE1*** | GTTCTGATAAGGCATCAGCCTCG |
| | CATGAGTAGGCTTCTAATAACGG |
| ***ACT7*** | TCCATGAAACAACTTACAACTCCATCA |
| | CATCGTACTCACTCTTTGAAATCCACA |
| **ChIP-qPCR** | |
| ***YUC8_I*** | CACCCTTCTTCAACAATCACCATCG |
| | CCGACCATTTTTGTGTGTTACTATAG |
| ***YUC8_II*** | CAACGCCACATGGGATCTCTTCTTC |
| | GAGAGAATCAATGGAAGTTGTATTG |
| ***YUC8_III*** | GAGAATATGTTTCGTTTGATGGATC |
| | CCGACGGTCCAGCTCCGACGATGAC |
| ***YUC8_IV*** | GTTCTCGTCGTTGGTTGTGGAAAC |
| | CATCACGTGAAGAGAGCTTCTCAC |
| ***YUC8_V*** | AACAACCCACGAAACGCTCAAGG |
| | CTCTTTTGTCAGTGATCTCTGTCC |
| ***YUC8_VI*** | TGTTTTGGCCCTGTTGAACAAATCC |
| | GGCTTTATGTCATAGTATAATTTCGC |
| ***SAUR20_I*** | GTGATGGAGACATGTTGTTCTGTTG |
| | CTTCTCCATGTTCTGTGATGATCC |
| ***SAUR20_II*** | GAGACAGAACCATGTTCTCTGTTC |
| | GATCAACTACTAATGAGTTGAAACC |
| ***SAUR20_III*** | AGCCTTCATTTCAAGCTCTTCTCAG |
| | GTCACATTGATGAATGTATCTTCAG |
| ***SAUR20_IV*** | ATGACCAAACCAATTGGAACTACTC |
| | GAGTCAGACGCGCTACCATTGCG |
| ***SAUR22_I*** | GAGATTCAAGAATAGATCATCACAG |
| | CACAAGGGCTTGTTTGCTCAATG |
| ***SAUR22_II*** | GAATCTTTTCATTCATCTTCAGATTTGC |
| | TTGCACCCAGTAGACTTCTCACC |
| ***SAUR22_III*** | TGCAACCGACACGACATGTCAATAC |
| | GGGAAACATATACTTTTCTAACCATG |
| ***SAUR22_IV*** | GTGCCATTGAGGCTTCCAGTAATC |
| | GAGATGTTGATCGCTACTTCTTGG |

| | |
|---|---|
| ***PRE1_I*** | TTACCCTCTACTTCTAAAATATGCAG |
| | TGTATGTGATCGTGAATACGTTTAC |
| ***PRE1_II*** | TCTCCCGTGACATCATTGTCGAG |
| | AAGAAATTGAGAATGTGATAAGAGAG |
| ***PRE1_III*** | TGCCACATTGTTGAACATGTCGAAC |
| | ACGGAGCTTAGATACGAGGTCAATC |
| ***PRE1_IV*** | TGTCGATGAAGATAGCCCTGAAG |
| | TAAGATTACATGGATAGGCTTGTC |
| ***PRE1_V*** | ACCCCATCTTAACTGTCACAAAAACG |
| | ACATGCATGTAGAATTATGAGATATC |
